# ABHD11 mediated deglutarylation regulates the TCA cycle and T cell metabolism

**DOI:** 10.1101/2024.09.11.612392

**Authors:** Guinevere L Grice, Eleanor Minogue, Hudson W Coates, Mekdes Debela, Nicole Kaneider-Kaser, P Robin Antrobus, Randall S Johnson, James A Nathan

**Affiliations:** Cambridge Institute of Therapeutic Immunology & Infectious Disease (CITIID), Jeffrey Cheah Biomedical Centre, Department of Medicine, University of Cambridge, Cambridge, CB2 0AW; Department of Physiology, Development and Neuroscience, University of Cambridge, Cambridge, CB2 3DY; Cambridge Institute for Medical Research, University of Cambridge, Cambridge; Department of Cell and Molecular Biology, Karolinska Institut, 171 77 Stockholm, Sweden

**Author notes:** Equal contributions (Guinevere L Grice, Eleanor Minogue, Hudson W Coates).

**Keywords:** ABHD11, α-ketoglutarate, 2-oxoglutarate, lipoic acid, glutarate, glutarylation, lipoylation, TCA cycle

## Abstract

Glutarate is an intermediate of amino acid catabolism and an important metabolite for reprogramming T cell immunity, exerting its effects by inhibition of histone demethylase enzymes or through glutarylation. However, how distinct glutarate modifications are regulated is unclear. Here, we uncover a deglutarylation pathway that couples amino acid catabolism to tricarboxylic acid (TCA) cycle function. By examining how glutarate can form conjugates with lipoate, an essential mitochondrial modification for the TCA cycle, we find that Alpha Beta Hydrolase Domain 11 (ABHD11) protects against the formation of glutaryl-lipoyl adducts. Mechanistically, ABHD11 acts as a thioesterase to selectively remove glutaryl adducts from lipoate, maintaining integrity of the TCA cycle. Functionally, ABHD11 influences the metabolic reprogramming of human T cells, increasing central memory T cell formation and attenuating oxidative phosphorylation. These results uncover ABHD11 as a selective deglutarylating enzyme and highlight that targeting ABHD11 offers a potential approach to metabolically reprogramme cytotoxic T cells.

## Introduction

Post translational modifications (PTMs) by small molecule metabolites are increasingly recognised as determinants of cellular metabolic reprogramming and cell fate decisions. Examples such as acetylation and succination highlight the importance of these modifications, with diverse cellular functions that range from chromatin regulation to immune effector responses^1–4^. Glutarate, a product of lysine or tryptophan catabolism, is a particularly interesting metabolite as it can be conjugated onto proteins or lipids (glutarylation), and metabolically inhibit 2-oxoglutarate (α-ketoglutarate) dependent dioxygenases (2-OGDDs) involved in epigenetic regulation^5–7^. Our understanding of the biology of glutarylation is in its infancy, but glutarate PTMs have important consequences for mitochondrial function^5,8,9^, metabolism^7^, and T cell immunity^6^.

Glutarate is formed by the conversion of 2-oxoadipic acid to glutaryl-coenzyme A (glutaryl-CoA) within the mitochondrion. Glutaryl-CoA can then generate free glutarate or be metabolised to acetyl-CoA. Human germline mutations in glutarate metabolism highlight its fundamental biological role, with loss of function mutations in glutaryl-CoA dehydrogenase (GCDH) resulting in glutarate accumulation and glutaric aciduria type 1 (GA1): a rare autosomal disorder characterised by dystonia, developmental delay and often death in early childhood^10^.

Glutarate conjugation to lysine residues (K_glu_) was first observed on mitochondrial proteins, with glutarylation of carbomoyl phosphate synthetase 1 (CPS1) reducing enzyme activity and altering ammonia clearance^5^. The regulatory nature of this PTM was supported by observations that dietary tryptophan supplementation alters K_glu_, and that SIRT5 can act as a K_glu_ deglutarylating enzyme^5,11^. Glutarylation has also been observed on histones, altering chromatin structure and dynamics^12^.

Recently, we uncovered that glutarylation is a dynamic modification in CD8+ T cells, with glutarylation patterns changing with T cell receptor (TCR) activation^6^. Glutarate accumulation resulted in K_glu_, as expected, but we also identified additional functions of glutarate. Firstly, it acts as a competitive inhibitor of 2-OGDD function, and secondly, glutarate accumulation impaired activity of the pyruvate dehydrogenase complex (PDHc). This latter finding is of particular interest, as rather than simply modifying lysine residues within PDH, glutarate formed a thioester with its lipoylated catalytic arm.

Lipoylation is an essential mitochondrial modification arising from conjugation of a redox-sensitive fatty acid (lipoic acid or lipoate) to defined lysine residues (K_Lp_)^13,14^. To date, five mitochondrial enzymes are known to require lipoylation for catalysis: the PDHc, 2-oxoglutarate dehydrogenase complex (OGDHc), 2-oxoadipate dehydrogenase complex (OADHc), branched-chain alpha-ketoacid dehydrogenase complex (BCKDHc), and the glycine cleavage system complex (GCVc). Lipoylation drives catalysis of these ketoacid dehydrogenase complexes via the cyclical oxidation and reduction of its thiol ring, described as the ‘swinging arm’ of the dehydrogenases. However, the reactive thiols of lipoate also render it to attack, resulting in lipoyl conjugates or adducts, such as we observed with glutarate accumulation^6^. Additionally, we and others have observed that other lipoyl adducts can be formed when lipid peroxidation products accumulate, or following nitrosylation during macrophage activation^15–18^, but how lipoate or lipoyl conjugates are regulated or removed remains poorly understood.

In this study we show that glutaryl-lipoyl adducts (Lp_glu_) are constantly formed on ketoacid dehydrogenases, and that ABHD11 is a selective glutaryl-lipoyl thioesterase that prevents the formation of these adducts, maintaining and preserving TCA cycle function. We additionally show that ABHD11 is regulated within immune cells, with cytotoxic T lymphocytes (CTLs) and pro-inflammatory Th1 cells expressing higher levels of ABHD11 than CD8+ memory T cells and anti-inflammatory Th2 cells. Inhibition of ABHD11 in CD8+ T cells increases the central memory T cell (T_CM_) pool and attenuates mitochondrial oxidative phosphorylation. These data illustrate the role of ABHD11 in regulating lipoylation and glutarylation, while also highlighting the therapeutic potential of targeting ABHD11 in CD8+ T cells to alter metabolism and increase stemness, a desire that is particularly pertinent in the field of immunotherapy.

## Results

### ABHD11 deficiency results in glutarylation of ketoacid dehydrogenases

Alpha Beta Hydrolase Domain 11 (ABHD11) is a mitochondrial hydrolase that maintains functional lipoylation^16^, but how this occurs and the nature of the lipoyl adducts formed was unclear. Given the identification of glutaryl-lipoyl adducts forming under glutarate accumulation^6^, we hypothesised that ABHD11 may reverse glutarylation of ketoacid dehydrogenases, maintaining enzymatic activity (**Fig. 1a**). We therefore explored whether ABHD11 inhibition or increased glutarylation resulted in similar lipoyl modifications.

**Fig. 1.**
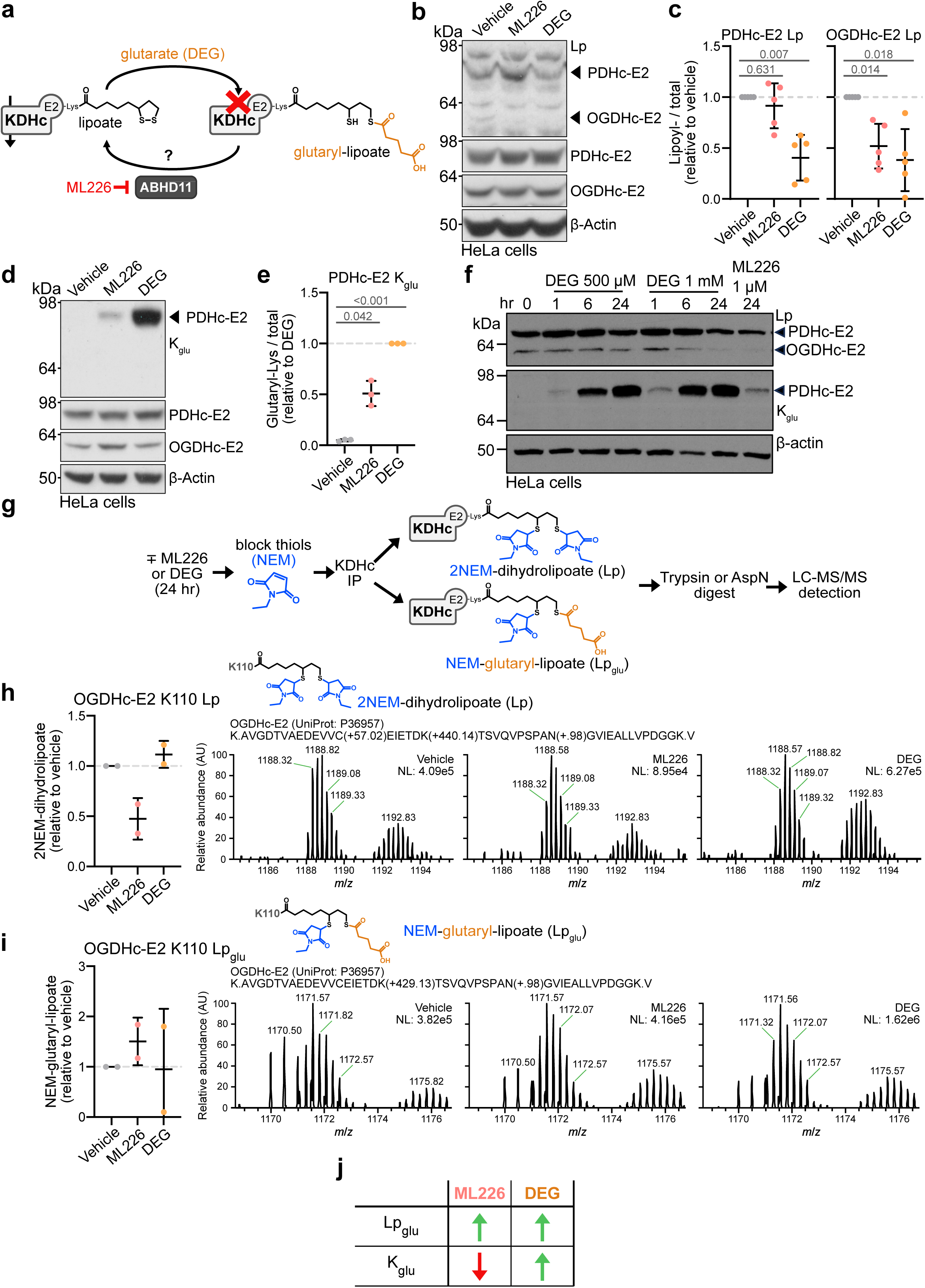
ABHD11 inhibition results in glutarylation of ketoacid dehydrogenases. **(a)** Glutaryl adduct formation on the lipoylated E2 subunit of ketoacid dehydrogenases (KDHc) and its putative removal by ABHD11. **(b-d)** Lipoyl (Lp) and glutaryl modifications on KDHc. HeLa cells were treated with 1 μM ML226 or 1 mM DEG for 24 hr and Lp modification of PDHc-E2 and OGDHc-E2 were determined using immunoblotting **(b)**. Visualisation of glutaryl modfications using a K_glu_ antibody **(d)** Quantification of Lp **(c)** or K_glu_ **(e).** *Mean ± SD; n = 3–5; one-way ANOVA and Dunnett’s post-hoc test.* **(f)** Time-course of the effect of DEG or ML226 treatment in HeLa cells on Lp or K_glu_ levels. **(g)** Schematic of LC-MS/MS analysis to detect Lp and Lp_glu_ modifications. **(h, i)** HeLa cells were treated with 2.5 μM ML226 or 1 mM DEG for 6 hr, or 1 μM ML226 or 1 mM DEG for 24 hr, OGDHc-E2 was immunoprecipitated, and K110 Lp **(h)** or K110 Lp_glu_ **(i)** was quantified using LC-MS/MS. Peptide abundance was normalised to the total abundance of modified peptides and adjusted relative to the vehicle condition. *Mean ± SD; n = 2*. Chromatograms are representative. **(j)** Differing consequences of ABHD11 inhibition and glutarate treatment.

We first determined if glutarylation of ketoacid dehydrogenases occurred following ABHD11 inhibition, similarly to that observed following treatment with cell-permeable glutarate (diethylglutarate, DEG)^6^. HeLa cells were treated with ML226, a selective inhibitor of ABHD11^16,19,20^, or DEG for 24 hr (**Fig. 1b-e**). Protein glutarylation was visualised using a pan K_glu_ antibody. Functional lipoylation was observed using a lipoate specific antibody, which recognises the unmodified lipoate conjugates on the two most abundant ketoacid dehydrogenases (OGDHc and PDHc) and not lipoyl adducts^16^. ABHD11 inhibition reduced lipoylation of the OGDHc-E2 (dihydrolipoyllysine-residue succinyltransferase, DLST), with no clear effect on the PDHc-E2 (dihydrolipoamide acetyltransferase, DLAT) (**Fig. 1b, c**), as previously reported^16^. However, DEG treatment reduced lipoylation of both the PDHc-E2 and OGDHc-E2 (**Fig 1b, c**). This global reduction in lipoylation with DEG treatment was associated with increased protein glutarylation (**Fig. 1d, e**). Additionally, we detected glutarylation in the ML226 treated cells (**Fig. 1d, e**), consistent with the potential involvement of ABHD11 in regulating ketoacid dehydrogenase glutarylation. However, DEG and ML226 demonstrated different dynamics of PDHc glutarylation with 1 μM ML226 resulting in lower levels of K_glu_ than 1 mM DEG over a time course of 6 to 24 hr (**Fig. 1f**).

Given that the pan-glutaryl antibody was raised against K_glu_, it was unlikely to be sensitive for detecting lipoyl modifications and may underestimate the involvement of ABHD11. Therefore, we used liquid chromatography mass spectrometry (LC-MS)^6,16^ to detect whether lipoyl glutarylation (Lp_glu_) or K_glu_ occurred following ABHD11 inhibition (**Fig. 1g**). We focused on the OGDHc-E2 as prior work indicates that ABHD11 preferentially regulates OGDHc lipoylation and function^16,21^. HeLa cells were treated with ML226 for 6 or 24 hr and endogenous OGDHc-E2 immunoprecipitated for analysis by LC-MS (**Fig. 1h, i; Extended Data Fig. 1a**). ML226 treatment decreased lipoylation of the OGDHc-E2 K110 residue relative to total OGDHc-E2 levels (**Fig. 1h**), consistent with prior observations^16^. Conversely, we detected increased levels of a K110 Lp_glu_ modification when ABHD11 was inhibited (**Fig. 1i**). In contrast, whilst DEG treatment increased OGDHc-E2 K_glu_, ML226 treatment reduced lysine glutarylation (**Extended Data Fig. 1b**), suggesting a more selective function of ABHD11.

To confirm the increase in Lp_glu_ was specific to ABHD11 function, we generated ABHD11 deficient HeLa cells using sgRNA (**Extended Data Fig. 1d**) and subjected the immunoprecipitated OGDHc-E2 to LC-MS. We also generated GCDH deficient cells, which blocks the downstream metabolism of glutaryl-CoA, providing a genetic route to potentially increase glutarate levels (**Extended Data Fig. 1c, d**), as occurs in GA1 disease^10^. SgRNA targeting β2 microglobulin (β2m) was used as a non-specific control. Higher K110 Lp_glu_ was detected in ABHD11 null cells compared to β2m, with a variable increase in K110 Lp_glu_ observed following GCDH depletion (**Extended Data Fig. 1e**). ABHD11 or GCDH knockout did not change K_glu_ (**Extended Data Fig. 1f**).

We also examined if Lp_glu_ or K_glu_ could be detected on the PDHc following ABHD11 inhibition, as suggested by the K_glu_ antibody (**Fig. 1b-d**). LC-MS analysis of the PDHc is more complex, given the two lipoylated sites on the PDHc-E2 subunit, and we were only able to reproducibly detect lipoyl moieties on K259. Despite this limitation, we still observed a relative reduction in lipoylation and a concomitant increase in Lp_glu_ (**Extended Data Fig. 2a, b**). K_glu_ levels increased following DEG treatment^6^, but were decreased by ML226 (**Extended Data Fig. 2c**), as observed with the OGDHc-E2. Therefore, while the K_glu_ antibody fortuitously detected a glutaryl signal following ML226 treatment, DEG and ABHD11 inhibition result in distinct glutaryl modifications. DEG treatment increased all glutaryl modifications, while ABHD11 inhibition results in Lp_glu_ with a concomitant decrease in K_glu_ (**Fig. 1j**).

### ABHD11 is a glutaryl-lipoyl thioesterase

The finding that Lp_glu_ increased following ABHD11 inhibition suggested that the serine hydrolase activity of ABHD11 regulated the lipoyl adduct. Therefore, we developed several *in vitro* assays to determine if ABHD11 removed glutaryl adducts from the lipoylated OGDHc-E2. First, we expressed and purified ABHD11-Flag in HEK23T cells and confirmed its hydrolase activity using a generic esterase substrate, p-nitrophenol acetate (**Fig. 2a, b**). We then measured whether purified ABHD11 could remove Lp_glu_ adducts from immunoprecipitated OGDHc, thereby restoring detection of lipoylation by immunoblot (**Fig. 2c**). This assay relies on a fortunate property of the lipoyl antibody, in that it only detects the functional lipoate moiety and not lipoyl adducts that form following NEM treatment or ABHD11 loss^16^. We generated OGDHc-E2 Lp_glu_ adducts by treating HeLa cells with ML226, and immunoprecipitated the OGDHc-E2. We then measured the levels of lipoate by immunoblot following short incubations with purified ABHD11-Flag at 37 °C. ABHD11 restored lipoylation after 15 min, with full restoration by 30 min (**Fig. 2d, e**). Similar findings were observed following ABHD11 depletion using sgRNA (**Fig. 2f, g**). Therefore, ABHD11 can reverse Lp_glu_.

**Fig. 2.**
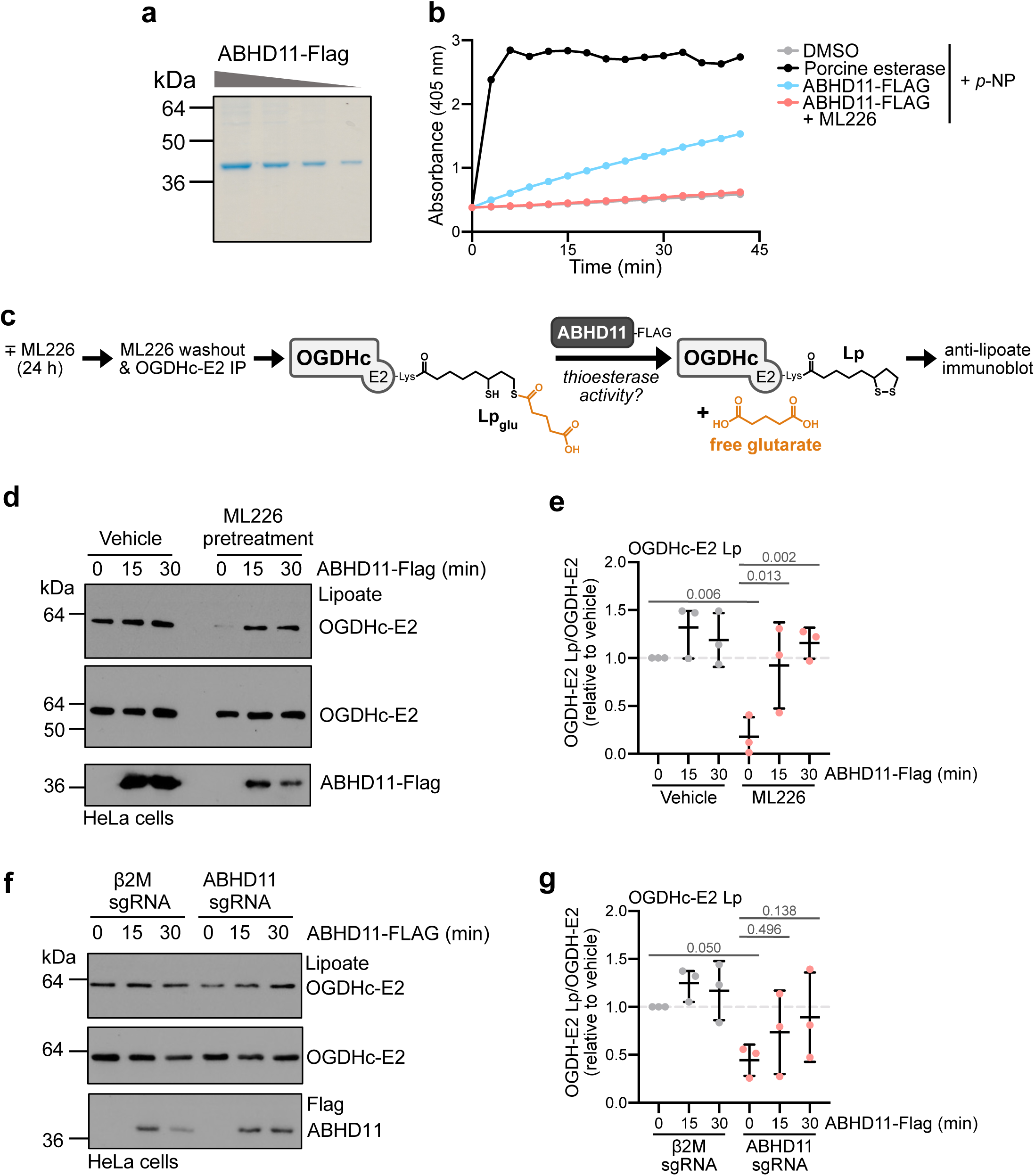
ABHD11 removes glutaryl adducts from lipoylated OGDHc-E2. **(a)** Representative Coomassie stain of purified ABHD11-Flag from HEK293T cells. **(b)** Generic esterase activity of ABHD11 using p-nitrophenyl acetate. Porcine esterase was used as a control. Representative assay of n = 3. **(c-g)** Purified ABHD11-Flag removes glutaryl-lipoate adducts from immunoprecipitated OGDHc-E2. Schematic of the experiments shown in **(c)**. Immunoprecipitated OGDHc-E2 from HeLa cells treated with 2.5 μM ML226 for 6 hr **(d, e)** or from ABHD11 deficient HeLa cells **(f, g)**. Immunoprecipitated OGDHc-E2 was incubated with ABHD11-Flag for the indicated times and lipoate levels measured by immunoblot **(d, f)** and quantified by normalisation to total OGDHc-E2 levels **(e, g)**. *n = 3; mean ± SD*, *Two-way ANOVA and Tukey’s post-hoc test*.

To confirm that ABHD11 acted directly on the Lp_glu_ adduct in an enzymatic manner, we used purified ABHD11 and measured its activity against a Lp_glu_ recombinant OGDHc-E2 peptide encompassing the minimal residues around the lipoylated K110 (**Extended Data Fig. 3a**). Thioesterase activity was measured by the addition of Ellman’s reagent (5,5′-dithiobis-(2-nitrobenzoic acid), DTNB) at 37 °C. This reagent provides a colorimetric readout when it binds to exposed free thiols thereby detecting the lipoate moiety if an adduct is removed (**Fig. 3a**). ABHD11 increased the availability of free thiols on the Lp_glu_ peptide compared to the vehicle control (**Fig. 3b, c**). ABHD11 had no effect on the unmodified peptide, nor a peptide with the lipoate thiol ring in an oxidised state (**Fig. 3b, c; Extended Data Fig. 3a**). The thioesterase activity of ABHD11 against the glutaryl moiety was dependent on enzyme and substrate concentration (**Extended Data Fig. 3b-f**). Moreover, ML226 treatment prevented the release of free thiols (**Fig. 3d, e**), demonstrating that the catalytic activity of ABHD11 was required.

**Fig. 3.**
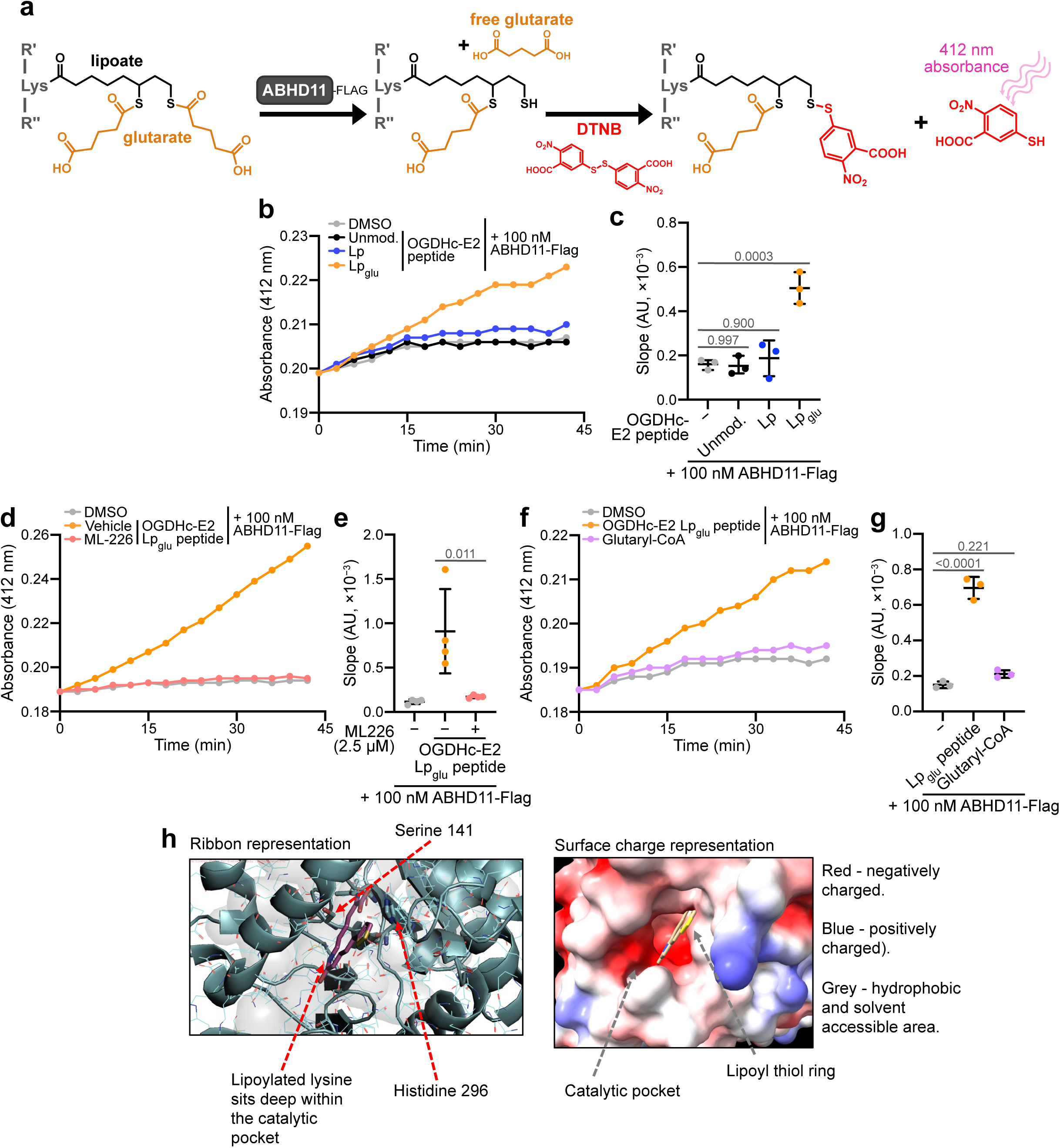
Glutaryl-lipoyl thioesterase activity of ABHD11. **(a)** Schematic of thioesterase assay using Ellman’s reagent (DTNB) to detect removal of glutaryl moieties from a synthetic OGDHc-E2 peptide (**Extended Data Fig. 3a**) by ABHD11-Flag. Any release of thiols by ABHD11 are detected via fluorescence at 412 nm following reaction with DTNB. **(b, c)** Thioesterase activity of ABHD11 (100 nM) on 100 μM of the Lp, Lp_glu_ and unmodified OGDHc-E2 peptide. *n = 3; mean ± SD*, *One-way ANOVA and Dunnett’s post-hoc test.* **(d, e)** Thioesterase activity of ABHD11 (100nM) against 100 μM Lp_glu_ OGDHc-E2 peptide with or without ML226 treatment (2.5μM). *n = 4; mean ± SD*, *One-way ANOVA and Tukey’s post-hoc test.* **(f, g)** Activity of ABHD11 (100 nM) on the OGDHc-E2 peptide (100 μM) in comparison to glutaryl-CoA (100 μM). *n = 3; mean ± SD*, *One-way ANOVA and Tukey’s post-hoc test.* **(h)** *In silico* modelling of ABHD11 to a marine α/β hydrolase fold esterase (RCSB.org ID:7c4d.1) (2.03 Å)) with lipoate moiety. Catalytic S141 and H296 indicated. Ribbon (left) and surface charge (right) models shown.

To explore the specificity of ABHD11 for glutaryl conjugates, we determined if ABHD11 could remove the glutaryl moiety from CoA using DTNB (**Fig. 3f, g**). However, unlike the glutaryl-lipoyl adduct, no free thiols were generated (**Fig. 3f, g**), indicating that ABHD11 acts as a selective glutaryl-lipoyl thioesterase. In support of these findings, we performed substrate docking modelling within the enzymatic cleft of ABHD11, based on the crystal structure of a related marine α/β hydrolase, with esterase activity^22^ (**Fig. 3h**). The lipoate moiety docked within the catalytic within the catalytic pocket, proximal to the catalytic serine-141 and histidine-296 residues (**Fig. 3h**). Together, these findings confirm that ABHD11 is a selective thioesterase that removes glutaryl moieties from the lipoate arm of the OGDHc.

### Glutaryl-lipoylation has distinct outcomes compared to protein glutarylation

Our enzymatic studies indicated ABHD11 has activity against Lp_glu_ but it was important to examine how ABHD11 inhibition functionally altered metabolism compared to perturbing cellular glutarate levels. Prior studies show that ABHD11 deficiency impairs the TCA cycle, and drives metabolic inhibition of 2-OGDDs through accumulation of 2-hydroxyglutarate (2-HG), activating the HIF response and altering the activity of some 2-OGDDs involved in modulating histone (e.g. lysine demethylases) and DNA methylation (e.g. TET enzymes)^16,21,23^ (**Fig. 4a**). Alternatively, DEG treatment has pleotropic effects on metabolism, altering Lp_glu_, K_glu_, and competitively inhibiting 2-OGDDs^6^. Therefore, we treated HeLa cells with either ML226 or DEG for 24 hr, and compared the downstream consequences on metabolism and 2-OGDD function (**Fig. 4a**).

**Fig. 4.**
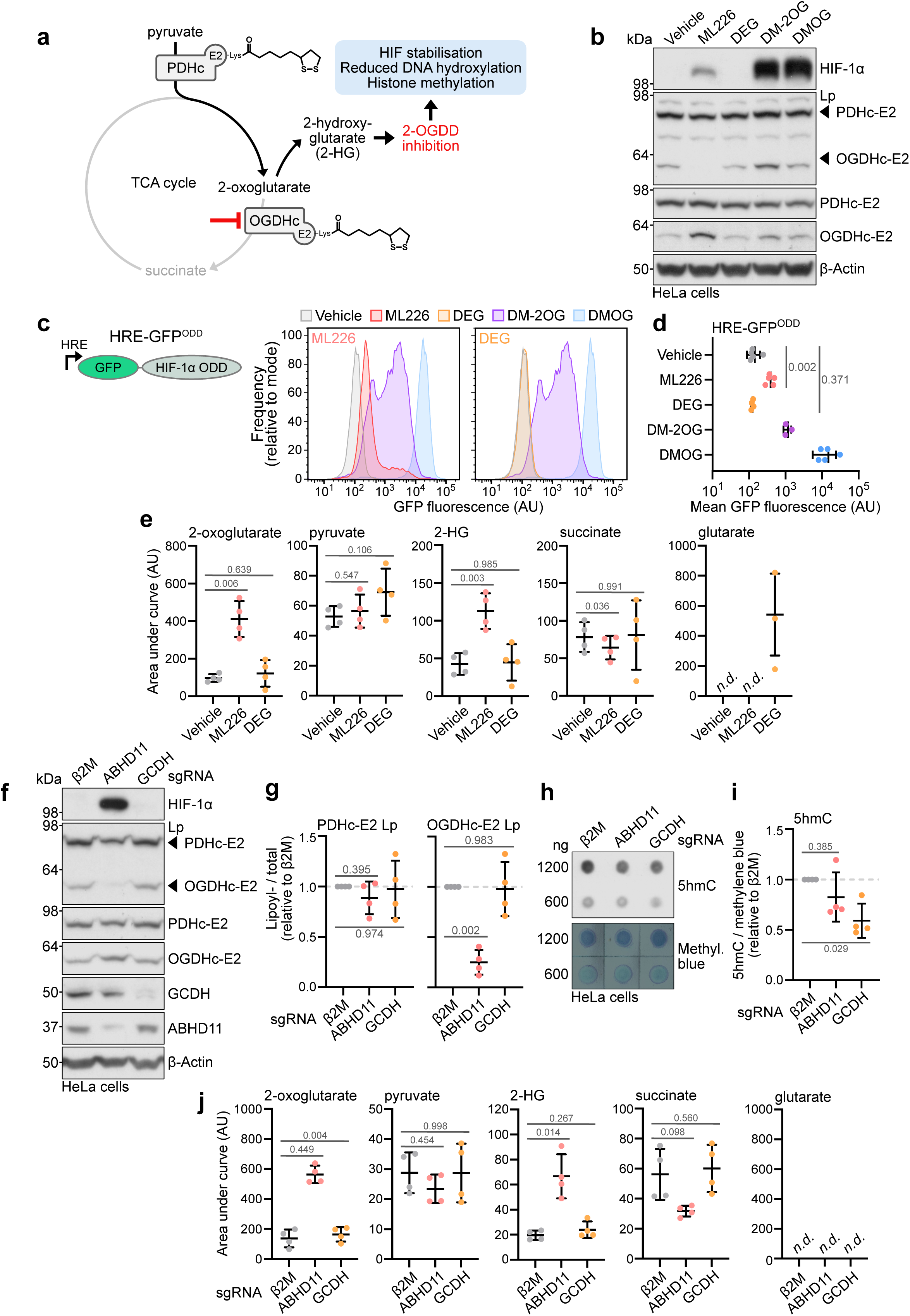
Glutaryl-lipoylation has distinct outcomes compared to protein glutarylation. **(a)** Schematic of outcomes of OGDHc inhibition on the TCA cycle and 2-OGDD function. 2-OG accumulation can drive L-2HG production and result in impaired 2-OGDD activity. Exogenous DM-2OG can also drive L-2HG accumulation and inhibit 2-OGDDs. DMOG directly inhibits 2-OGDDs. **(b-d)** Comparison of ML226 to DEG, DM-2OG treatment on HIF-1α stabilisation and activity. HeLa cells were treated with 2.5 μM ML226, 1 mM DEG, 6 mM DM-2OG for 6 hr. DMOG (1 mM, 6 hr) was used as a control. HIF-1α levels were determined by immunoblotting. *Representative of n = 3*. **(c, d)** HeLa HRE-GFP^ODD^ cells were treated with 1 μM ML226 or 1 mM DEG for 24 hr and GFP levels were determined using flow cytometry. For visualisation purposes, ML226 and DEG conditions are plotted separately. Quantification of mean GFP fluorescence **(d)**. *Mean ± SD; n = 3–5; one-way ANOVA and Dunnett’s post-hoc test.* **(e)** Metabolite analysis of HeLa cells treated with 1 μM ML226 or 1 mM DEG for 24 hr by LC-MS/MS. *Mean ± SD; n = 4; one-way ANOVA and Dunnett’s post-hoc test; n.d., not detected.* **(f-g)** Comparison of HIF-1α levels in Cas9-expressing HeLa cells transduced with sgRNAs targeting β2M, ABHD11 or GCDH. After 11 days, HIF-1α and Lp modifications were determined using immunoblotting. β-actin served as a loading control. Quantification of Lp normalised to total PDHc-E2 or OGDHc-E2 and adjusted relative to the β2M condition **(g)**. *Mean ± SD; n = 4; one-way ANOVA and Dunnett’s post-hoc test*. **(h, i)** 5hmC levels in ABHD11 or GCDH deficient cells **(f)**. DNA was extracted 11 days post sgRNA transduction. 5hmC levels were determined using dot blotting and quantified by normalising to total DNA staining (methylene blue) and adjusted relative to the β2M condition **(i)**. *Mean ± SD; n = 4; one-way ANOVA and Dunnett’s post-hoc test.* **(j)** Metabolite analysis of ABHD11 or GCDH deficient HeLa cells **(f)**. The indicated metabolites were quantified using LC-MS/MS. *Mean ± SD; n = 4; one-way ANOVA and Dunnett’s post-hoc test; n.d., not detected*.

ML226 treatment resulted in HIF-1α stabilisation, upregulation of HIF-1 target genes (*CAIX* and *VEGF*), and activation of a HIF-1 fluorescent reporter (HRE-GFP^ODD^) in HeLa cells, similarly to treatment with cell-permeable 2-OG (dimethyl 2-OG, DM-2OG) that simulates impaired OGDHc function and 2-hydroxyglutarate accumulation (2-HG)^23,24^ (**Fig. 4b-d; Extended Data Fig, 4a**). However, DEG treatment did not alter HIF-1α stabilisation nor activity (**Fig. 4b-d; Extended Data Fig. 4a**). We also observed differences in the levels of 5-hydroxymethylcytosine (5hmC) and histone H3 lysine methylation marks between ML226 and DEG treatment (**Extended Data Fig. 4b-e**). These findings suggested that DEG did not drive 2-HG formation (and particularly the L enantiomeric form), as occurs with OGDHc inhibition^23,24^. Indeed, when we undertook LC-MS analysis of TCA cycle metabolites, ML226 treatment resulted in 2-OG and 2-HG accumulation but this did not occur following DEG treatment (**Fig. 4e**).

Next, we compared the phenotypic consequences for ABHD11 loss with GCDH depletion. ABHD11 loss increased HIF-1α levels and decreased OGDHC-E2 lipoylation, similarly to ML226 inhibition, but no changes in HIF-1α levels or lipoylation were observed in the GCDH KO cells (**Fig. 4f, g**). Although 5hmC levels were reduced in both ABHD11 and GCDH KO cells (**Fig. 4h, i**), levels of H3K4-, H3K9-, or H3K27-trimethylation were unchanged (**Extended Data Fig. 4f, g**). The preservation of lipoylation following GCDH loss was surprising, given the findings with DEG and LC-MS analysis. However, when we measured cellular glutarate levels in the GCDH KO cells, they were below the level of detection (**Fig. 4j**), suggesting that metabolic adaptation can occur to prevent glutarate accumulation, potentially missing transient glutaryl modifications. However, ABHD11 KO cells had increased 2-OG and 2-HG levels, indicating that accumulation of Lp_glu_ predominantly alters OGDHc activity. Thus, ABHD11-mediated regulation of Lp_glu_ is distinct from other perturbations of glutarate metabolism, and predominantly alters OGDHc activity.

### ABHD11 influences human CD8+ T cell differentiation

Given the diverse roles for glutarylation in CD8^+^ T cell fate and function^6^, we wanted to interrogate the contribution of Lp_glu_ and ABHD11 function. We first examined ABHD11 expression in published hematopoietic cell proteomes^25^ to find which immune cells express high levels of ABHD11, and under what conditions ABHD11 is regulated. ABHD11 was expressed at low levels in naive CD4+ and CD8+ T cells but markedly accumulated upon T cell receptor activation in both cell types (**Fig. 5a**). Amongst activated CD4+ T cell populations, Th1 cells contained higher levels of ABHD11 than Th2 cells, and both Th17 and iTreg cells contained low levels of ABHD11 akin to the naive cell. Memory CD8+ T cells expressed less ABHD11 than cytotoxic CD8+ T cells, although its levels were higher than any CD4+ T cell population. Levels of ABHD11 were low in natural killer cells and eosinophils, but higher in mast cells. These cell type and activation-dependent changes in ABHD11 expression strongly suggested it has a functional role in T cell biology, and particularly CD8+ T cells. Therefore, we determined if ABHD11 inhibition disrupts lipoylation in CD8+ T cells, similarly to HeLa cells. CD8^+^ T cells were isolated from healthy human donor peripheral blood mononuclear cells, activated using CD3/CD28 beads, and continuously cultured with ML226 for 4 days. Immunoblotting revealed a reduction in OGDHc-E2 lipoylation but no change in PDHc-E2 lipoylation was detected (**Fig. 5b, c**). Therefore, ABHD11 predominantly alters OGDHc function, as observed in the HeLa cells.

**Fig. 5.**
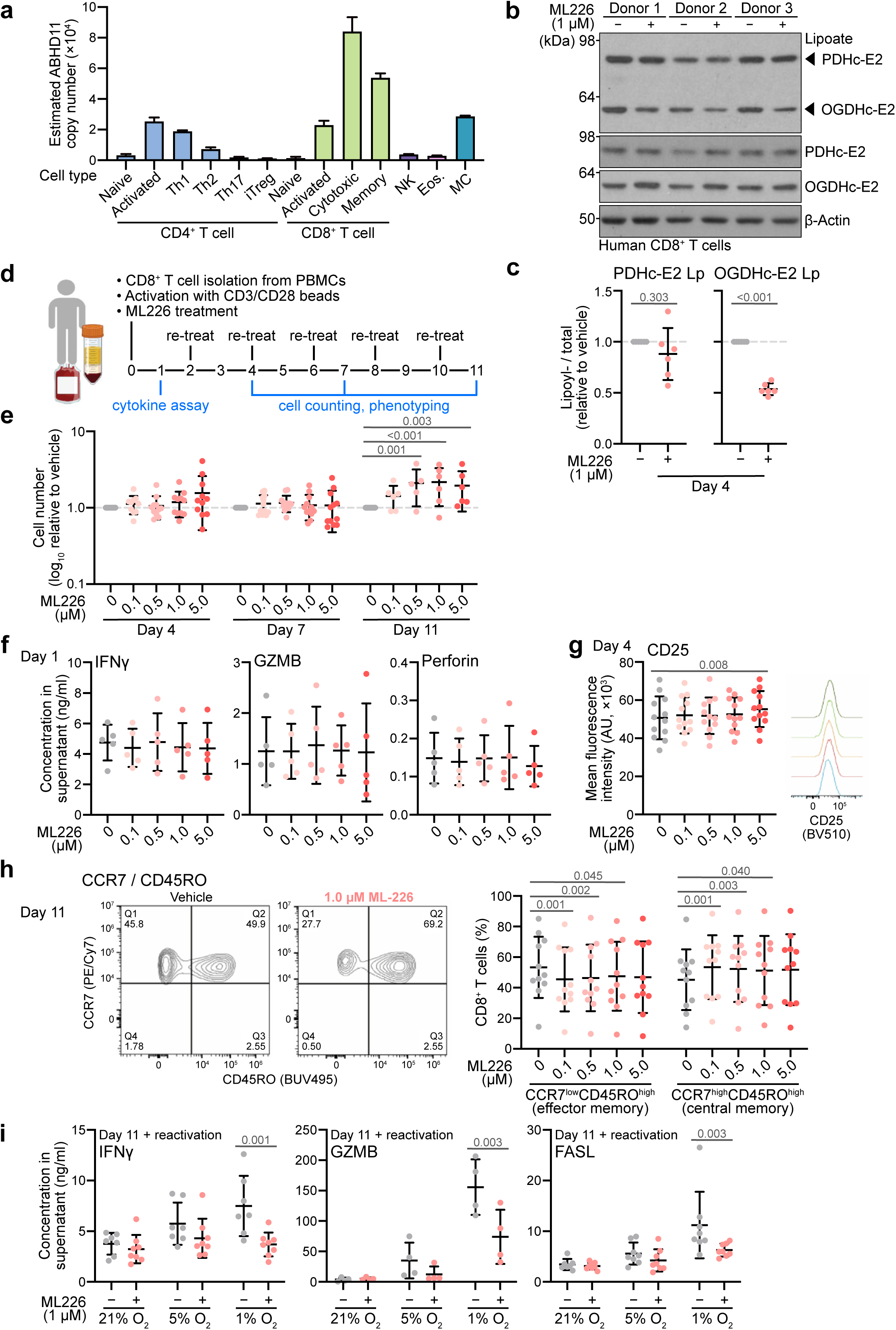
ABHD11 influences human CD8+ T cell differentiation. **(a)** Protein copy number of ABHD11 in hematopoietic cell proteomes from the ImmPRes database. *NK, natural killer; Eos., eosinophil; MC, mast cell.* **(b, c)** CD8^+^ T cells were isolated from healthy donor peripheral blood mononuclear cells, activated with CD3/CD28 beads, and continuously treated with 1 μM ML226 for 4 days. Lp modifications on PDHc-E2 and OGDHc-E2 were determined using immunoblotting (representative of three donors. Quantification of Lp **(c)**, normalised to total PHDc-E2 or OGDHc-E2 and adjusted relative to the vehicle condition. Each data point represents one donor. *Mean ± SD; n = 6; paired two-tailed t-test.* **(d)** Timeline of human CD8^+^ T cell isolation, treatment, and phenotypic assays for experiments **(e-h)**. CD8^+^ T cells were isolated from healthy donor peripheral blood mononuclear cells, activated with CD3/CD28 beads, and continuously treated with 1 μM ML226 for up to 11 days. **(e)** CD8+ T cell numbers following activation and continuous culture with the indicated concentrations of ML226 for 4, 7, or 11 days. Numbers are relative to vehicle control for each timepoint. *Mean ± SD; n = 12 for Day 4 and Day 7; n = 6 for Day 11; two-way ANOVA and Dunnett’s post-hoc test.* **(f)** Cytokine levels in cell culture supernatants from **(d)** 1 day post-activation. Cytokine levels were determined using flow cytometry. *Mean ± SD; n = 5; one-way ANOVA and Dunnett’s post-hoc test.* **(g)** CD25 levels in CD8+ T cells 4 days post-activation **(d)**, determined using flow cytometry. *Mean ± SD; n = 5; one-way ANOVA and Dunnett’s post-hoc test. Flow cytogram is representative.* **(h)** Percentage of CCR7 and CD45RO-expressing cells 11 days post-activation were determined using flow cytometry. *Mean ± SD; n = 6; one-way ANOVA and Dunnett’s post-hoc test per subpopulation. Flow cytogram is representative.* **(i)** CD8+ T cells were acclimatised to the indicated oxygen concentrations for 2 hr, activated with CD3/CD28 beads, and continuously cultured at the indicated oxygen concentrations with 1 mM ML226 for 11 days. Cells were then re-activated for 4 hr and cytokine levels in cell culture supernatant were determined using flow cytometry. *Mean ± SD; n = 4–8; two-way ANOVA and Dunnett’s post-hoc test*.

We next examined the role of ABHD11 in CD8+ T cell fate and function, and cultured human CD8^+^ T cells continuously with ML226 treatment for up to 11 days (**Fig. 5d**). ML226 was well tolerated and did not affect cell growth during short-term culture (4 or 7 days), but cell numbers were increased after 11 days (**Fig. 5e**). Short term treatment with ML226 did not alter T cell receptor activation, as there was no change in the secretion of key cytokines (**Fig. 5f; Extended Data Fig. 5a**) nor the expression of CD25, a marker of activation (**Fig. 5g**). There were generally no significant changes in T cell exhaustion markers, except at the highest dose of ML226 at day 11 (**Extended Data Fig. 5b, c**). However, we did observe that ML226 drove a memory T cell phenotype, as after 11 days treatment, there was an increase in the CCR7^hi^CD45RO^hi^ central memory T cell population (T_CM_), with a corresponding decrease in the CCR7^lo^CD45RO^hi^ effector memory T cell population (T_EM_) (**Fig. 5h; Extended Data Fig. 5d**).

To further explore the functional role of ABHD11, we reactivated CD8+ T cells at different oxygen tensions, including those more representative of physiological oxygen levels. Cytokine release was not affected following reactivation of CD8+ T cells on day 11 at 21 %, or 5 % oxygen, but when reactivation was performed at 1% oxygen, levels akin to the tumour microenvironment (TME) or sites of infection, ML226 treated CD8+ T cells showed reduced secretion of interferon gamma (IFNy), granzyme B (GZMB), and Fas ligand (FASL) (**Fig 5i; Extended Data Fig. 5e**). TNF-α, perforin, granulysin, and granzyme A (GZMA) showed a downward trend in 1% oxygen but this was not significant when compared to vehicle treated cells (**Extended Data Fig. 5e**). Together, these findings support a role for ABHD11 in controlling CD8+ T cell differentiation, with implications for the cytotoxic activity that occurs in low oxygen environments.

### ABHD11 regulation of the OGDHc alters CD8+ T cell metabolism

Lastly, to determine how ABHD11 inhibition increases CD8+ T_CM_ populations, we examined the metabolic consequences of this treatment and the effect on histone marks regulated by L-2HG (S-2HG). We quantified total H3K4me3, H3K9me3, and H3K27me3 in CD8+ T cells, following 4 days of culture at 21%, 5%, and 1% oxygen. ML226 did not alter the abundance of histone methylation marks at 21% or 5% oxygen, but in 1% oxygen, reduced levels of H3K4me3 and H3K9me3 were observed (**Fig. 6a**). Surprisingly, no changes in 5hmC levels were observed (**Fig. 6b, c**), in contrast to the changes seen in HeLa cells. These data indicate that metabolic inhibition of 2-OGDD activity via 2-HG was unlikely to be responsible for the phenotypic changes in T cells following ABHD11 inhibition with ML226, and distinct from prior observations with L-2HG^26^. Therefore, we focused on the metabolic consequences of ABHD11 inhibition in CD8+ T cells, and the implications for OGDHc inactivation.

**Fig. 6.**
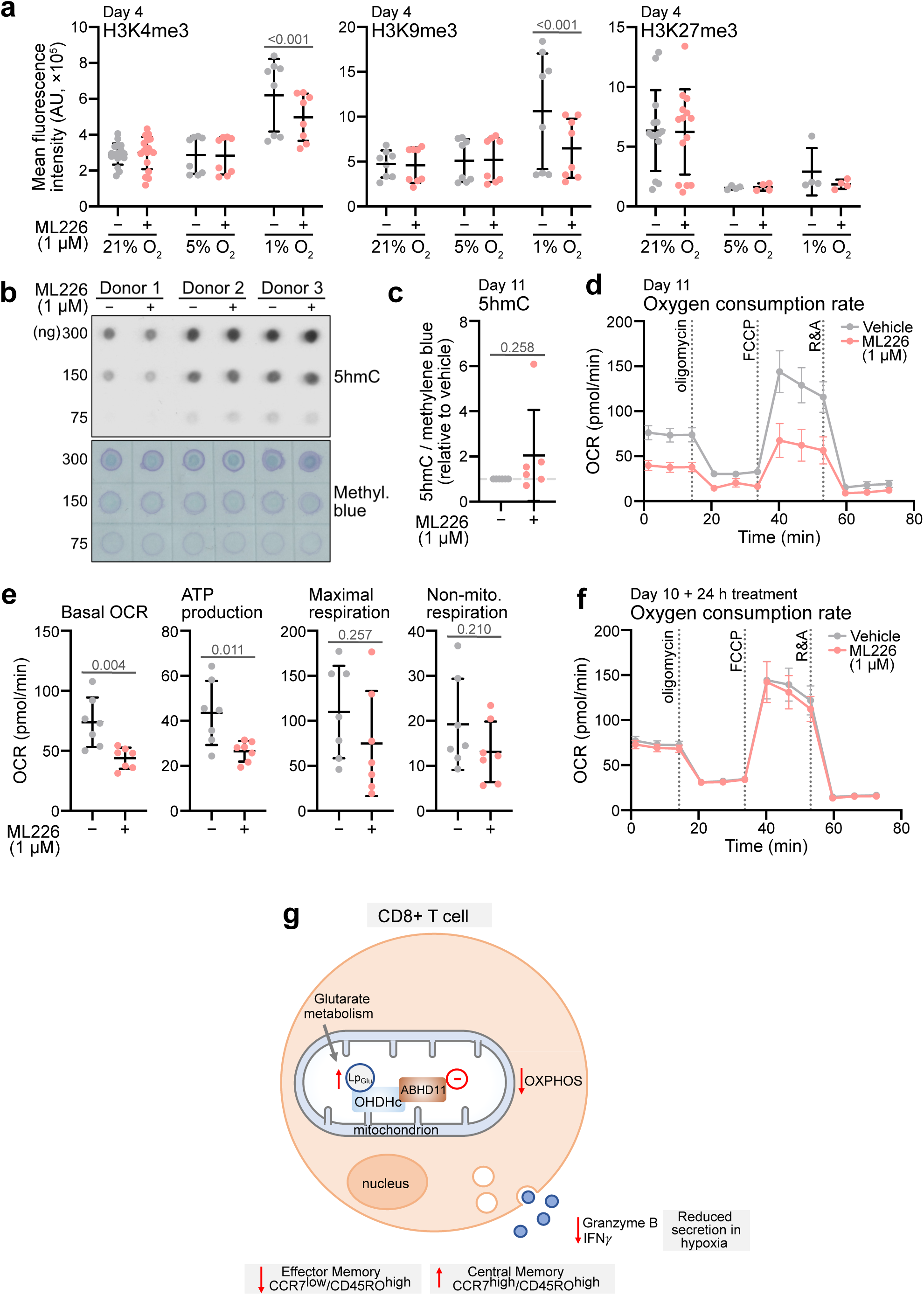
Effect of ABHD11 inhibition on CD8+ T cell metabolism. **(a)** CD8+ T cells were isolated from healthy donor peripheral blood mononuclear cells, acclimatised to the indicated oxygen concentrations for 2 hr, activated with CD3/CD28 beads, and continuously cultured at the indicated oxygen concentrations plus 1 mM ML226 for 4 days. H3K4me3, H3K9me3, and H3K27me3 levels were determined using flow cytometry. *Mean ± SD; n = 8–18; two-way ANOVA and Dunnett’s post-hoc test.* **(b, c)** Activated CD8+ T cells were continuously treated with 1 mM ML226 for 11 days and 5hmC levels were determined using dot blotting. Quantification of 5hmC levels **(c)**, normalised to total DNA staining by methylene blue and adjusted relative to the vehicle condition. *Mean ± SD; n = 6; mean ± SD; n = 6; paired two-tailed t-test.* **(d, e)** CD8+ T cells were isolated from healthy donor peripheral blood mononuclear cells, activated with CD3/CD28 beads, and continuously treated with 1 mM ML226 for 11 days. Oxygen consumption rate (OCR) during a mitochondrial stress test (1 μM oligomycin, 1.5 μM FCCP, 100 nM rotenone plus 1 μM antimycin) was quantified using a Seahorse XFe96 Extracellular Flux Analyzer. Quantification of basal OCR, ATP production, maximal respiration, and non-mitochondrial respiration in **(e)**. *Mean ± SEM **(d)** or SD **(e)**; n = 6; unpaired two-tailed t-test.* **(f)** Activated CD8^+^ T cells were cultured for 10 days and treated with 1 mM ML226 for 24 hours. OCR during a mitochondrial stress test was quantified using a Seahorse XFe96 Extracellular Flux Analyzer. *Mean ± SEM; n = 4*. **(g)** Schematic of effect of decreased ABHD11 function on CD8+ T cell phenotype and function. Loss or decreased ABHD11 reduced OXPHOS via Lp_glu_ accumulation on the OGDHc-E2. Decreased TCA cycle function and OXPHOS is associated with a phenotypic switch to central memory CD8+ T cells, and reduced secretion of IFNγ and Granzyme B under hypoxic conditions.

ML226 treatment in primary CD8+ T cells reduced the basal oxygen consumption rate (OCR), similarly to the loss of OGDHc function observed in HeLa ABHD11 KO studies (**Fig. 6d, e**). However, there were no significant changes in extracellular acidification (ECAR) or glycolysis (measured by a standard glycolysis stress test) in CD8+ T cells treated with ML226 (**Extended Data Fig. 6a-e**), indicating again that ML226 has more of an effect on OGDHc activity than PDHc activity. Prolonged inhibition of ABHD11 was required for the changes in basal OCR, as ML226 treatment for 24 hr at day 10 did not alter OCR or ECAR (**Fig. 6f; Extended Data Fig. 6f**). These findings are consistent with CD8+ T cell metabolism being intrinsically linked to function, and as memory CD8+ T cells are known to be less metabolically active than CTLs^27^, the reduced ABHD11 expression that occurs when CTLs transition to memory cells (**Fig. 5a**) may contribute to metabolic differences between these two cell states (**Fig. 6g**).

## Discussion

Lipoylation and glutarylation are both recognised as dynamic PTMs involved in mitochondrial metabolism, cell fate determination, and immune regulation, but the enzymatic regulation of these modifications is only just being appreciated^5,6,12–16,21,23,28–34^. By distinguishing between lipid and protein glutarylation we were able to uncover a role for ABHD11 in removing Lp_glu_ adducts, thereby maintaining activity of essential ketoacid dehydrogenases. Our studies highlight the selective regulation of an acyl-lipoyl conjugate (thioester bond), which is distinct from the role of Sirtuins (SIRTs) in cleaving acyl modifications from lysine residues (amide bonds). SIRT4 has reported lipoamidase activity^28^, while SIRT5 was the first described K_glu_ deglutarylating enzyme^5^. However, both SIRT4 and SIRT5 have more widespread activity in removing other K_acyl_ chains and regulating other mitochondrial enzymes^35–39^. In contrast, ABHD11 does not so far demonstrate amidase activity, and has a predominant effect on Lp_glu_ through hydrolysis of the thioester bond.

The preferential regulation of the OGDHc-E2 by ABHD11 may relate to the number of K_Lp_ residues compared to the PHDc-E2. Both the OGDHc and PDHc-E2 require K_Lp_ for catalytic activity but the PDHc-E2 may be protected from lipoyl adducts, as it contains two K_Lp_ residues. A further explanation may relate to the same E2 subunit being shared between the OGDHc and OADHc. Lp_glu_ is an intermediate formed by the OADHc that coverts 2-oxoadipate to glutaryl-CoA, potentially accounting for the susceptibility of this E2 to glutaryl modifications. However, persistent glutarylation of the OGDHc-E2 would prevent the formation of glutaryl-CoA by the OADHc, and also render the OGDHc inactive. Interestingly, not only did we not observe an accumulation of free glutarate when ABHD11 was depleted, but we also observed a decrease in K_glu_ on the OGDHc-E2, suggesting that ABHD11 activity may alter the available glutaryl-CoA pool. Testing this by detecting glutaryl-CoA by LC-MS has not been possible, and further enzymology is required to understand the relationship between the OGDHc, OADHc, and glutaryl-CoA. However, our data does implicate Lp_glu_ in a negative feedback loop to control glutaryl-CoA metabolism.

*In silico* modelling of ABHD11 is consistent with the lipoyl domain docking within the catalytic pocket. We previously showed that the serine-141 and histidine-296 are required for enzymatic activity^16^, and docking predicts that the thiol ring of lipoate is proximal to these residues. Other factors that drive ABHD11 substrate specificity remain to be determined, and will require further structural studies. However, it is noteworthy that ABHD11 had no activity against the glutaryl-CoA thioester, suggesting that either CoA cannot dock within the catalytic pocket, or that the OGDHc-E2 peptide guides specificity by binding to the relatively charged surface of the catalytic site. Interestingly, we did not detect changes in other Lp_acyl_ modifications on the OGDHc-E2^16^, but it is plausible that ABHD11 may remove other acyl chains, and these additional functions of ABHD11 may relate to the phenotype observed in ABHD11 null mice^40^.

The role of ABHD11 against Lp_glu_ adducts provides further evidence for the regulatory nature of lipoylation. Whilst essential for maintaining key metabolic pathways, ours and other studies demonstrate that lipoyl adducts can serve roles more akin to signalling, as demonstrated for other metabolite PTMs. Lipoyl adducts formed by changes in oxidative stress or catabolic stress can alter TCA cycle function, allowing the cell to adapt to these insults, as demonstrated for transient lipoyl nitrosylation during macrophage activation^15,32^. Lipoyl adducts are also implicated in a recently described cell death pathway, cuproptosis^41^. We now show that Lp_glu_ adducts also influence T cell function, with impaired 2-OG metabolism associated with increasing the pool of CD8+ T_CM_ cells. Therefore, the regulation of these different lipoyl modifications provides opportunities to both alter immune cell function and potentially manipulate cuproptosis. Understanding how these different lipoyl modifications interact will be a necessary focus of future studies.

High levels of ABHD11 expression in immune cell populations^25^ highlight its fundamental role in these cells, with likely broad implications for immune effector responses. ABHD11 inhibition increases the pool of CD8+ T_CM_ cells: a T cell population that displays stem cell like properties, with enhanced activity against pathogens and cancer cells compared to T_EM_ cells^27^. We also observed that in 1% oxygen, ABHD11 activity altered the secretion of cytokines involved in CD8+ T cell cytotoxic activity. Further studies will focus on understanding the full extent of manipulating ABHD11 on anti-tumour activity and viral infections, but our findings are highly relevant to the cancer immunotherapy field, where increasing CD8+ T_CM_ would be beneficial for chimeric antigen receptor (CAR) T therapy^42^. Additionally, our data suggest that ABHD11 and the regulation of lipoyl adducts may influence the function and activity of other immune effector cells, including CD4+ T cells and macrophages. Therefore, ABHD11-mediated regulation of Lp_glu_ modifications is not only central for the TCA cycle but likely to influence several immune cell phenotypes, highlighting the potential clinical utility of our work.

## Methods

### Cell lines and reagents

HEK293T and HeLa cells were maintained in high-glucose DMEM (Sigma-Aldrich D6429) containing 10% fetal calf serum (Sigma-Aldrich P4333), 100 U/ml penicillin, and 100 µg/ml streptomycin at 37 °C and 5% CO_2_. Clonal HeLa cell lines expressing Cas9 (with a hygromycin selection cassette) or an HRE-GFP^ODD^ reporter construct were previously generated by the laboratory^23^. Low-oxygen incubations (5% or 1% O_2_) were carried out in a Whitley H35 Hypoxystation. All cell lines were authenticated and routinely tested for mycoplasma. The major reagents used in this study are listed in **Supplementary Data 1**.

### CD8+ T cell isolation and culture

Human peripheral blood mononuclear cells (PBMCs) were obtained from the National Health Service (NHS) Blood and Transplant (NHSBT) (Addenbrooke’s Hospital, Cambridge, UK). Ethical approval was obtained from the East of England-Cambridge Central Research Ethics Committee (06/Q0108/281) and consent was obtained from all participants.

PBMCs from healthy donors were isolated from blood using discontinuous plasma Percoll gradients. Total CD8+ T cells were purified from PBMCs by magnetic-activated cell sorting using CD8 microbeads (Miltenyi 130-045-201). Immediately after isolation, CD8+ T cells were activated with anti-human CD3/CD28 Dynabeads (Thermo Fisher Scientific 11131D) at a 1:1 cell:bead ratio and cultured in complete RPMI 1640 supplemented with 10% FBS, 100 U/ml penicillin, 100 µg/ml streptomycin, and 30 U/ml IL-2 (Sigma-Aldrich 11147528001). For experiments involving oxygen control, purified CD8^+^ T cells were immediately acclimatized to the required oxygen tension (21%, 5%, or 1% O_2_) for 2 hr in complete RPMI 1640 supplemented with 10% FBS, 100 U/ml penicillin, and 100 µg/ml streptomycin, prior to activation with CD3/CD28 Dynabeads and 30 U/ml IL-2.

For experiments, CD8+ T cells were seeded at a density of 5×10^5^ cells/ml medium. Treatments with ML226 (or 0.1% v/v DMSO vehicle) were performed at CD3/CD28 activation unless otherwise indicated. Low-oxygen incubations (5% or 1% O_2_) were carried out in a Ruskinn SCI-tive workstation. Cell number was determined using a CellDrop Automated Cell Counter (DeNovix).

### Plasmids

CRISPR/Cas9 sgRNA vectors targeting β2M and ABHD11 were previously generated by the laboratory^16^. Sequences targeting GCDH (RefSeq ID: 2639) were identified using VBC-Score and the top three hits were cloned into the pKLV-U6sgRNA pGK Puro-2A-BFP vector (Addgene #50946) using BbsI cloning. Plasmid identity was verified using Sanger sequencing and the most efficient sgRNA (as determined by anti-GCDH immunoblotting) was used in subsequent experiments. sgRNA sequences are listed in **Supplementary Data 1**.

### Lentivirus preparation and transduction

To prepare lentivirus particles, 4×10^5^ HEK293T cells were seeded into 6-well plates and transfected the next day with 0.66 µg expression plasmid, 0.44 µg pCMV-dR8.91 (gag/pol), and 0.88 µg pMD.G (VSVG) using 8 µl FuGENE HD transfection reagent (Promega). After 48 hr, viral supernatants were collected, filtered (0.45 µm), and stored at –80 °C in single-use aliquots.

For lentiviral transduction, 0.2×10^6^ HeLa-Cas9 cells were seeded into 24-well plates containing 500 µl viral supernatant. After 24 hr, cells were expanded into 6-well dishes and selected in 1 µg/ml puromycin for 3 days, then further expanded in non-selective medium. Experiments were performed using mixed populations 11 days post-transduction.

### Immunoblotting

To profile protein lipoylation and lysine glutarylation, 0.5×10^6^ HeLa cells or 1.0×10^6^ T cells were lysed in SDS loading buffer (2% [w/v] SDS, 50 mM Tris [pH 7.4], 150 mM NaCl, 1 mM dithiothreitol, 10% [v/v] glycerol, and 0.2% [v/v] benzonase nuclease [Sigma-Aldrich E1014]) and heated at 90 °C for 5 min. For T cell analysis, 1×10^6^ cells were washed with ice-cold PBS and lysed on ice in 60 µL RIPA buffer (Thermo Fisher Scientific 89900) supplemented with Halt Protease Inhibitor Cocktail (100x, Thermo Fisher Scientific 78429).

Samples were separated on duplicate SDS-PAGE gels, transferred to PVDF membranes, and blocked in 2% fatty acid free bovine serum albumin (BSA) and 5% skim milk powder in PBS containing 0.2% Tween-20 (PBST) for lipoate detection, or 5% skim milk powder in PBST for glutaryl-lysine detection. Blots were incubated in the appropriate primary antibodies (**Supplementary Data 1**) followed by HRP-conjugated secondary antibody in blocking buffer. Immunoblots were developed on film using HRP-conjugated secondary antibody and Pierce ECL Western Blotting Substrate (Thermo Fisher Scientific 32106). To quantify total PDHc-E2 and OGDHc-E2, lipoate blots were stripped using Restore Western Blot Stripping Buffer (Thermo Fisher 21059), re-blocked in 5% skim milk in PBST, and re-immunoblotted using appropriate primary antibodies. The signal was quantified using Image Studio Lite v5.5 (LI-COR Biosciences). HIF-1α, ABHD11, GCDH were analysed by re-immunoblotting glutaryl-lysine blots using appropriate primary antibodies.

To profile histone methylation marks, immunoblotting was performed as described above using 12% SDS-PAGE gels and blocking in 5% skim milk in PBST. Membranes were stripped and re-blocked between each primary antibody incubation. Methylation marks were normalised to total H3 levels, then adjusted relative to the control condition.

### Quantitative PCR

To analyse mRNA expression, total RNA was harvested from cells using the PureLink RNA Mini Kit (Thermo Fisher) and 1 µg was reverse transcribed using the ProtoScript II First Strand cDNA Synthesis Kit with oligo-dT primers and murine RNase inhibitor (New England Biolabs). Quantitative PCR was performed in technical triplicate using 20 ng cDNA, 1× Power SYBR Green PCR Master Mix (Thermo Fisher), and 0.5 µM forward and reverse primers (**Supplementary Data 1**), using an ABI 7900HT Real-Time PCR system (Applied Biosystems). Target gene expression was calculated using Quantstudio v1.3 (Applied Biosystems), normalised to the *ACTB* housekeeping gene using the ΔΔC_T_ method, and adjusted relative to the control condition.

### Flow cytometry

To profile the stability of the HeLa HRE-GFP^ODD^ cells, live cells were washed twice with cold PBS and immediately analysed using a LSR II flow cytometer (BD Biosciences). GFP signal was detected by a 488 nm laser with 530/30 filter. Data analysis were performed using FlowJo v10.9 (BD Biosciences).

To quantify protein levels in CD8+ T cells, single-cell suspensions were stained using the LIVE/DEAD Fixable Near-IR Dead Cell Stain Kit (Thermo Fisher Scientific 10119), followed by surface and intracellular staining with fluorochrome-labelled antibodies (**Supplementary Data 1**). Staining of cytoplasmic and nuclear antigens was performed using the Cytofix/Cytoperm Fixation/Permeabilization Kit (BD Biosciences 554714) and Transcription Factor Buffer Set (BD Biosciences 562725), respectively. Data were collected using an Aurora flow cytometer (Cytek Biosciences).

### DNA hydroxymethylation analysis

5hmC levels were quantified as previously described^16^. Briefly, genomic DNA was harvested from 1×10^6^ cells using the Puregene Cell Kit (Qiagen). DNA was diluted to 100 ng/µl in 0.4 M NaOH and 10 mM EDTA (pH 8.0) and denatured by heating at 100 °C for 10 min. Serially diluted DNA was spotted in 3×2 µl droplets onto a HyBond NX nylon membrane (Amersham) and allowed to dry. Membranes were incubated in 2× SSC buffer (Sigma-Aldrich J60839.K2), air-dried, and UV-crosslinked at 120,000 µJ/cm^2^ for 150 s using a UV Stratalinker 2400 (Stratagene). For immunoblotting, membranes were blocked in 5% skim milk powder and 1% BSA in PBST for 1 hr and incubated in anti-5-hydroxymethylcytosine antibody. Signal was developed on film using HRP-conjugated secondary antibody and SuperSignal West Dura Extended Duration Substrate (Thermo Fisher Scientific 34075). For total DNA staining, membranes were incubated in 0.1% (w/v) methylene blue solution containing 0.5 M sodium acetate (pH 5.2) overnight at room temperature. After 3×5 min washes with distilled water, membranes were air-dried and scanned. Signal from 5-hydroxymethylcytosine immunoblots was quantified using Image Studio Lite v5.5 (LI-COR Biosciences) or Image J, and normalised to methylene blue signal, then adjusted relative to the control condition. Within each biological replicate, adjusted data were averaged amongst the serial dilutions of genomic DNA.

### Immunoprecipitation

To enrich PDHc-E2 or OGDHc-E2 for proteomics analysis, 5×10^7^ cells were lysed in 1.5 ml IP buffer (50 mM HEPES [pH 7.0], 150 mM NaCl, 1% [v/v] IGEPAL CA-630, 1 mM phenylmethylsulfonyl fluoride, 10 mM tris[2-carboxyethyl]phosphine hydrochloride, 100 mM nicotinamide, 1× Roche cOmplete protease inhibitor cocktail) and rotated at 4 °C for 30 min. Lysates were pelleted at 16,900 *g* and 4 °C for 10 min, and 5% of the supernatant volume was collected as an input control. The remaining supernatant was mixed with 10 mM N-ethylmaleimide, rotated at 4 °C for 1 hr, and pre-cleared with Pierce Protein G magnetic beads (Thermo Fisher Scientific 88848) by rotating at 4 °C for 1 hr. The supernatant was then mixed with 100 µl Protein G magnetic beads bound to 10 µl anti-PDHc-E2 or anti-OGDHc-E2 antibody and immunoprecipitated at 4 °C for 16 hr. Beads were washed 4× with IP wash buffer (50 mM Tris-HCl [pH 7.4], 150 mM NaCl, 0.1% [v/v] IGEPAL CA-630), then 3× with IP wash buffer lacking IGEPAL CA-630. Proteins were eluted by heating in SDS loading buffer at 90 °C for 5 min, and lipoylation phenotypes were verified by immunoblotting the eluate alongside the input control. The above procedure was scaled down for *in vitro* assays requiring smaller-scale immunoprecipitations.

### Post-translational modification analysis

For mass spectrometry analysis of PDHc-E2 or OGDHc-E2 modifications, immunoprecipitation eluates were separated on a pre-cast 4−12% NuPAGE gel (Thermo Fisher Scientific NP0321BOX) and protein bands were stained using SimplyBlue SafeStain solution (Thermo Fisher Scientific LC6060). After several washes with distilled water, bands corresponding to PDHc-E2 (∼75 kDa) or OGDHc-E2 (∼55 kDa) were excised using a sterile scalpel and transferred to Protein LoBind tubes (Eppendorf).

Mass Spectrometry data (LC-MS/MS) was acquired as described previously^6^. An Orbitrap Fusion Lumos coupled to an Ultimate 3000 RSLC nano UHPLC equipped with a 100 µm ID x 2 cm Acclaim PepMap Precolumn (Thermo Fisher Scientific) and a 75 µm ID x 50 cm, 2 µm particle Acclaim PepMap RSLC analytical column was used. Loading solvent was 0.1% FA with analytical solvents A: 0.1% FA and B: 80% MeCN + 0.1% FA. Samples were loaded at 5 µl/minute loading solvent for 5 min before beginning the analytical gradient. The analytical gradient was 3-40% B over 42 min rising to 95% B by 45 min followed by a 4 min wash at 95% B and equilibration at 3% solvent B for 10 min. Columns were held at 40°C. Data was acquired in a DDA fashion with the following settings: MS1: 380-1500 Th, 120,000 resolution, 4×10^5^ AGC target, 50 ms maximum injection time. MS2: Quadrupole isolation at an isolation width of m/z 1.6, HCD fragmentation (NCE 30) with fragment ions scanning in the Orbitrap from m/z 110, 5×10^4^AGC target, 100 ms maximum injection time. Dynamic exclusion was set to +/− 10 ppm for 60 s. MS2 fragmentation was trigged on precursors 5×10^4^ counts and above.

Raw files were processed using PEAKS Studio (version 8.0, Bioinformatics Solutions Inc.) Variable modifications at PEAKS DB stage: oxidation, carbamidomethylatation, lipoylation, 1X and 2x NEM lipoylation, Glutaryl lipoate, NEM Glutaryl lipoate and 309 built in modifications at PEAKS PTM stage. The area under the curve (AOC) for each peptide were extracted from PEAKS peptide list, with AOCs being calculated by PEAKS. All raw data is included in **Supplemental Data 2**.

### ABHD11 in silico modelling

#### Homology model building

The 2.03 A X-ray structure of marine esterase (Alph/Beta hydrolase fold, RCSB.org: 7c4d.1, www.pdb.org)^22^ was downloaded and homology similarity of ABHD11 assessed in Swiss-model software (proMod3 3.3.0, www.swiss.org), in the server platform where the ABHD11 model was initially built. To refine the new output (model ABHD11) building locally, a Coot^43^ command line and interactive molecular graphics interface software were used. The 7c4d.1(reference) and ABHD11 coordinates, similarity, and active site pocket residues were topologically overlaid onto corresponding positions in the reference coordinate and visualised in Coot^43^. The ABHD11 coordinate (ABHD11.mmcif) was then able to be manually edited, and super-imposed over the reference electron density map (.mtz file of 7c4d.1). The structure, active site residues, outliers of loops, side chains, or unreliable atomic coordinates were closely monitored. Particular attention was paid to a poorly resolved area of 50 amino acids at the n-terminus, which was assigned to zero occupancy in the newly generated model.

#### Preparing the ligand coordinate

The (xx(lipoyl)lysine) ligand was sketched manually using molview.org for visualisation and the ligand atoms were then converted into a crystallographic information file (.cif), atomic coordinate format. The hydrogen bond distances were monitored/corrected in SHELX^44^ and the output file was then read on Coot.

#### Ligand docking in the active site of ABHD11

Both the (xx(lipoly)lysine).cif and ABHD11.cif coordinates were submitted to the HADDOCK2.4 server^45^, where molecular docking for this experiment was performed based on 3D structures and ligand input. An output of 10 binding models from the system were released, based on different configurations and ligand coordinate interactions. The top scoring complex coordinate output file was selected based on the atomic interaction, atomic distance/ clashing, binding interface with ligand (xx(lipoyl)lysine) and surface charges in the active site pocket following display of each coordinate in Coot.

All 3D model figures, surface rendering and active site occupancy representations of the complex were created with PyMOL Version 3.0.4 and Pymol-plungin/command line script.

### ABHD11 purification

To purify ABHD11-Flag protein, 1 x 10^8^ HeLa cells expressing ABHD11-Flag were lysed in 1ml lysis buffer (50mM Tris-HCl pH7.4, 150mM NaCl, 1% Triton-×100, 1× Roche cOmplete protease inhibitor cocktail) and rotated at 4 °C for 1 hr. Lysates were pelleted at 16,900 *g* and 4 °C for 10 min, and 5% of the supernatant volume was collected as an input control. Remaining supernatants were then precleared with CL4B resin and rotated at 4 °C for 1 h. CL4B resins were pelleted at 16,900 *g* and 4 °C for 1 min and the supernatants were then incubated with 200 µl M2 magnetic beads (Sigma M8823) and immunoprecipitated at 4 °C for 16h. Beads were then washed four times with wash buffer (50mM Tris-HCl pH7.4, 150mM NaCl, 0.1% Triton-×100) and then twice with wash buffer lacking Triton-×100. ABHD11-Flag protein was then eluted in 600 µl 100 µg/ml 3X Flag peptide and rotated at 4 °C for 1 h. Elution was then repeated. Concentration and purity of ABHD11-Flag protein was then verified in eluates using Coomassie Simply Safe stain. Dependent on protein purity, pooled eluates were either dialysed immediately or further purified using anion exchange chromatography.

Purified ABHD11-Flag was dialysed against one litre of TBS (50mM Tris-HCl pH7.4, 150mM NaCl) with a magnetic stirrer 4 °C for 16h, followed by a second dialysis with a magnetic stirrer at 4 °C for 2 hr. Purity of ABHD11-Flag protein was then verified in the dialysed sample using Coomassie Simply Safe stain and the concentration compared with BSA standards. Dialysed ABHD11-Flag was then concentrated if required using a centrifugal filter column (Vivaspin 2, 10,000 MWCO) at 3,000 rpm and 4 °C for 3 min. After adding 10% glycerol, concentration and purity of ABHD11-Flag protein was verified by Coomassie Simply Safe stain and BSA standards. Aliquots were stored at −80°C.

### Thioesterase assays

#### ABHD11 reconstitution assay

1 x 10^6^ HeLa cells treated with or without 2.5 μM ML226 for 6 hr were lysed and immunoprecipitated as described. The OGDHc-E2 bound resins were washed in IP wash buffer (without IGEPAL CA-630) and resuspended in 70 μl TBS (50 mM Tris pH 7.4, 150mM NaCl). A 10μl aliquot of the resins was retained to verify the immunoprecipitation by immunoblot. For the assay, 20 μl resins were aliquoted into 3 × 1.5 ml eppendorfs for each condition, with 20 μl ABHD11-Flag or vehicle control added. Samples were incubated for 15 or 30 min at 37°C on a shaking heat block. At each endpoint samples were placed on ice, centrifuged at 3,000 rpm 4°C for 30 s and 4μl 6 × SDS loading buffer was added. Samples were then heated at 90°C for 5 min, centrifuged at 3,000 rpm for 30 s, placed on a magnet for 1 min, and supernatants were collected for SDS-PAGE analysis. Lipoate and OGDHc-E2 levels were measured by immunoblot.

#### DTNB assay

10 mg of unmodified OGDHc E2, lipoyl OGDHc E2 or diglutaryl lipoyl OGDHc E2 peptides were generated by Peptide Synthetics to > 90 % purity and reconstituted to 6.56 mM in DMSO. Glutaryl CoA (Glutaryl-CoA Salt, G9510-5MG, Sigma) was reconstituted in DMSO to 6.56 mM. Peptides or glutaryl CoA were subsequently diluted to 2.5mM in assay buffer (50 mM Tris-HCl pH 7.4, 100 mM NaCl). 5,5′-dithiobis-(2-nitrobenzoic acid) (DTNB) (Sigma) was dissolved in 10× PBS (pH 7.4) to make a 4 mg/ml (10 mM) solution and subsequently diluted in assay buffer to a final concentration of 0.2 mM. 100 μM of the peptides or glutaryl CoA were added to 100 nM ABHD11-Flag (50 mM Tris-HCl pH 7.4, 100 mM NaCl) and 0.2 mM DTNB reagent in a 96 well plate on ice. 2.5 μM ML226 (Cayman Chemicals) was added at the start of the reaction to experiments measuring ABHD11 inhibition. Absorbance was measured at 412 nm in a plate reader (CLARIOstar Plus Microplate, BMG LabTech) using kinetic mode with measurements every 45 s for 45 min at 37° C. Michaelis-Menten kinetics were calculated by first obtaining the concentration of TNB formed under the reactions (*c = A/bE*), where *c* is the concentration, *A* is absorbance, *b* is the pathlength (1 cm) and *E* is the molar absorptivity (molar extinction coefficient of TNB is 14,150 M^−1^cm^−1^)^46^. The velocity (*V, μM/min*) was calculate from the slope of the kinetic assays. The Michaelis-Menten model was plotted in GraphPad Prism and the *K_M_*, *k_cat_*, and *k_cat_/K_M_* calculated.

### Cytokine release assays

To quantify CD8+ T cell cytokine release, 200 µL of cell culture supernatant was collected 4– 24 hr post-activation, centrifuged at 2,000 *g* for 2 min to remove remaining cells, and stored at −20 °C until analysis. Samples were thawed at room temperature and cytokine concentrations (IL-2, IL-4, IL-6, IL-10, IL-17A, IFN-γ, TNF-α, soluble Fas, soluble FasL, granzyme A, granzyme B, perforin, and granulysin) was quantified using the LEGENDplex Human CD8/NK V02 Panel (Biolegend 741187), according to the manufacturer’s instructions. Data were collected using an Aurora flow cytometer (Cytek Biosciences) and analyzed using the LEGENDplex Data Analysis Software Suite (Qognit). All samples were analyzed in at least duplicate.

### Metabolic analysis

To measure oxygen consumption and extracellular acidification, a Seahorse XFe96 Extracellular Flux Analyzer (Agilent) was used. Here, 1.5×10^5^ CD8+ T cells were plated in Seahorse XF cell culture plates coated with poly-D-lysine (Sigma-Aldrich P7280) and preincubated in appropriate medium at 37°C and the absence of CO_2_ for 1 hr prior to treatments. Three basal measurements were made at 3 min intervals, followed by three measurements per treatment. A minimum of four technical replicates were analysed per biological replicate.

For mitochondrial stress tests, the preincubation medium consisted of Seahorse XF RPMI medium (pH 7.4; Agilent 10376-100) supplemented with 2 mM glutamine (Thermo Fisher Scientific 25030081) and 10 mM glucose (Thermo Fisher Scientific A2494001). Treatments comprised 1 μM oligomycin (Sigma-Aldrich 75351), 1.5 μM FCCP (Sigma-Aldrich C2920), and 100 nM rotenone (Sigma-Aldrich R8875) plus 1 μM antimycin A (Sigma-Aldrich A8674). For glycolysis stress tests, the preincubation medium consisted of Seahorse XF RPMI medium (pH 7.4) supplemented with 2 mM glutamine. Treatments comprised 10 mM glucose, 1 μM oligomycin, and 0 mM 2-deoxy-D-glucose (BioVision B1048-100).

### Metabolite quantification

1×10^6^ treated Hela cells or 1×10^6^ Cas9-expressing HeLa cells were harvested, washed with cold PBS, and metabolic activity quenched by freezing samples in dry ice and ethanol, and stored at −80°C. Before extraction, 10 µL of the internal standard solution (d_4_-Succinic acid, ^13^C_4_-Glutaric acid and ^13^C_5_-2-hydroxyglutarate; 100 μg / mL each) was added to all samples. Metabolites were subsequently extracted by addition of 600 μl ice-cold LC-MS grade methanol to the cell pellets, followed by 30 min sonication (Sweep mode). During sonication, ice was added to the ultrasound bath to maintain the temperature below 20° C. Following sonication, samples were centrifuged (12000 *g*, 15 min, 10°C) and 200 µL of the supernatant were transferred to LC-MS vials for the subsequent analysis.

Targeted analyses were performed on a Waters Xevo TQ-Sµ Triple Quadrupole Mass Spectrometer (Waters, Milford, MA). An Atlantis Premier BEH Z-HILIC FIT column (2.1 mm × 100 mm, 1.7 μm), equipped with an integrated guard column (Vanguard FIT), was used for the separation, with aqueous mobile phase consisting of 10 mM Ammonium Acetate pH = 9.1 (adjusted with NH4OH) in double-deionised water and organic mobile phase consisting of 0.1% formic acid in acetonitrile. Column oven was set at 30° C, and 1 µL was injected for each sample. The flow rate was set at 0.45 mL/min. For the separation, the following chromatographic gradient was used:

| Minute | %B | Flow rate (µl/min) |
| --- | --- | --- |
| 0.0 | 85 | 450 |
| 3.0 | 79 | 450 |
| 5.5 | 60 | 450 |
| 7.0 | 50 | 450 |
| 8.5 | 50 | 450 |
| 8.7 | 85 | 450 |
| 12.5 | 85 | 450 |

MS analyses were performed using electrospray ionisation (ESI) operating in the negative mode ionisation using the following parameters: capillary potential (1.0 kV), source temperature 150° C, and desolvation temperature 600°C. Compound identification was based on retention time and, at least, one single monitoring reaction (SRM) transition SRM matching the standards used to build the calibration curve. The retention times (RT) and the quantifier (Quant) and qualifier (Qual) SRM transitions for the reported compounds were as follows: 2-hydroxyglutarate (RT = 5.5 min, Quant: 147 > 57; Qual: 147 > 129); 2-oxoglutarate (RT = 5.4 min, Quant: 145 > 57); Glutarate (RT = 5.5 min, Quant: 131 > 69; Qual: 131 > 113); Pyruvate (RT = 1.7; Quant: 87 >43; Qual: 87 > 32) and Succinate (RT = 5.4 min; Quant: 117 > 73; Qual: 117 > 99). Raw data is included in **Supplemental Data 3**.

### Quantification and statistical analysis

Quantitative data are expressed as the mean of biological repeats ± 1 standard deviation (SD) or ± 1 standard error of the mean (SEM). P-values were calculated using two-tailed Student’s t-tests, or analysis of variance (ANOVA), as indicated in figure legends. Statistical analyses of the mass spectrometry data are described in the relevant method sections. The number of biologically independent repeats are specified in figure legends. Figures were prepared and statistical analyses performed using GraphPad Prism or R. Software used in this study is indicated in **Supplementary Data 1**.

### Data availability

Data from the lipoylation proteomics is shown in **Supplementary Data 2**. Data files from the metabolomics experiments is included in **Supplementary Data 3**.

### Code availability

No custom software or code was generated for these studies.

## Acknowledgements

We thank all members of the Nathan and Johnson labs for their helpful comments on the work and manuscript. The authors gratefully acknowledge Dr Antonio Checa and the Karolinska Institute Small Molecule Mass Spectrometry Core Facility (KI-SMMS), supported by KI/SLL, for support in the sample analyses. We also gratefully acknowledge the support of the Cambridge Institute of Medical Research Mass Spectrometry Facility, the flow cytometry facility of the School of the Biological Sciences of the University of Cambridge. Bioenergetic experiments were performed at the Medical Research Council Toxicology Unit, University of Cambridge, Cambridge, UK with thanks to Prof. Marion MacFarlane for access. This work was also supported by the NIHR BRC flow cytometry facility. This work was funded by a Wellcome Senior Clinical Research Fellowship to JAN (215477/Z/19/Z), a Wellcome Principal Research Fellowship (214283/Z/18/Z) to RSJ, and a Lister Institute Research Fellowship to JAN.

## Author contributions

Conceptualization, GLG, EM, RSJ, and JAN; Methodology, GLG, EM, HWC, MD, PRA, and JAN; Investigation, GLG, EM, HMW, PRA, MD and JAN; Writing – original draft, GLG, EM, HWC and JAN; Writing – reviewing and editing, all authors; Funding acquisition, RSJ, JAN; Resources, PRA, NK, RSJ and JAN; Supervision, NK, RSJ and JAN.

---

**Extended Data Fig 1.**
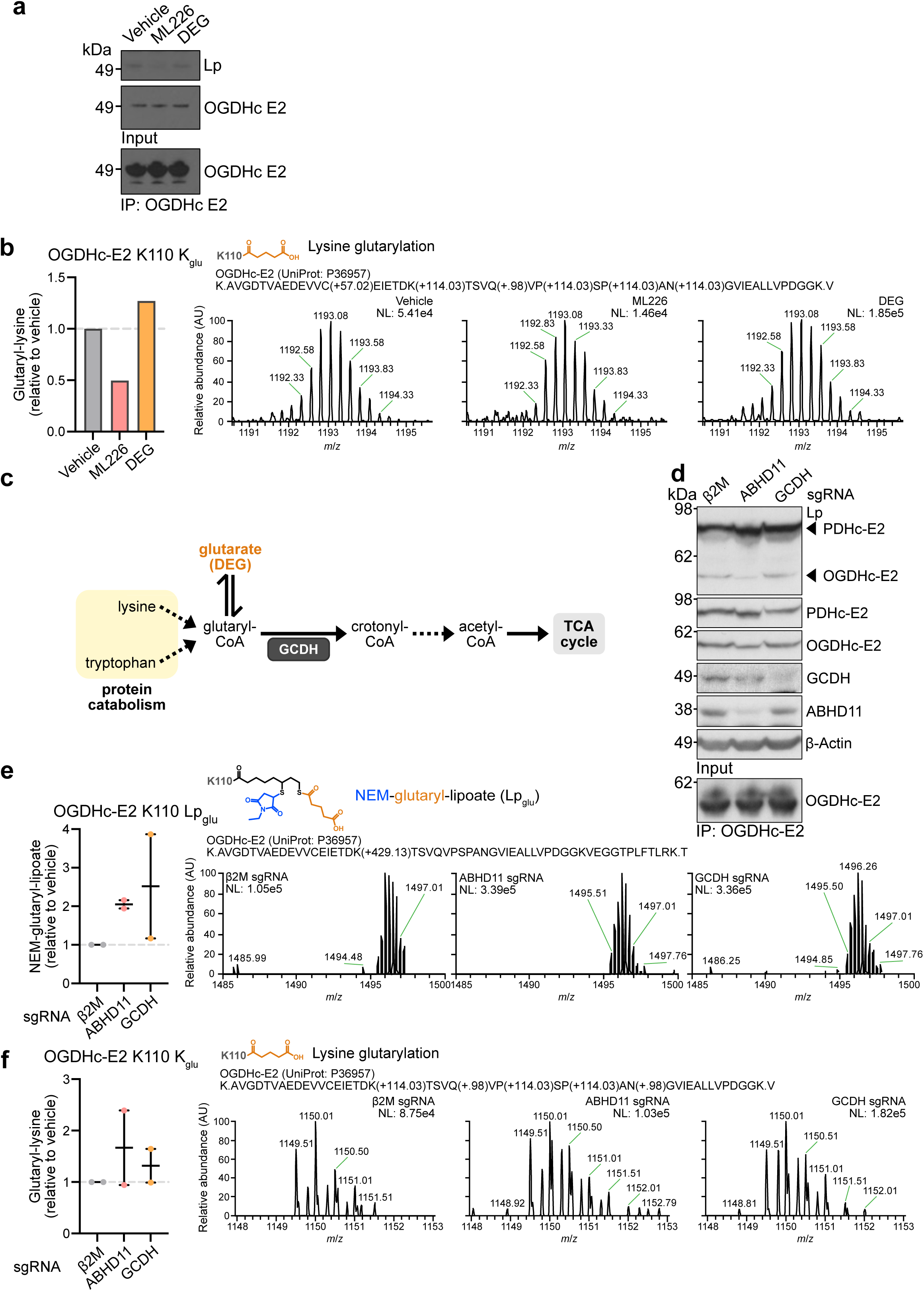
OGDHc glutaryl-lipoate adducts accumulate following ABHD11 depletion. **(a)** Representative OGDHc-E2 immunoprecipitation in HeLa cells for LC-MS/MS analysis of lipoate modifications. *n = 2*. **(b)** Quantification of OGDHc-E2 K110 lysine glutarylation (K_glu_) in cells from **Fig. 1h–i**. Peptide abundance was normalised to the total abundance of modified peptides and adjusted relative to the vehicle condition. Chromatograms are representative. **(c)** Schematic of glutaryl-CoA metabolism indicating action of GCDH. **(d)** Effect of ABHD11 or GCDH depletion on lipoylation. Cas9-expressing HeLa cells were transduced with sgRNAs targeting β2m, ABHD11 or GCDH for 11 days. Immunoblot is representative of lipoylation phenotype and OGDHc-E2 immunoprecipitation. **(e, f)** Quantification of OGDHc-E2 K110 Lp_glu_ **(e)** and K110 K_glu_ **(f)** in cells from **(d)**. Peptide abundance is normalised to the total abundance of modified peptides and adjusted relative to the β2M condition. Chromatograms are representative. *n = 2; mean ± SD*.

**Extended Data Fig. 2.**
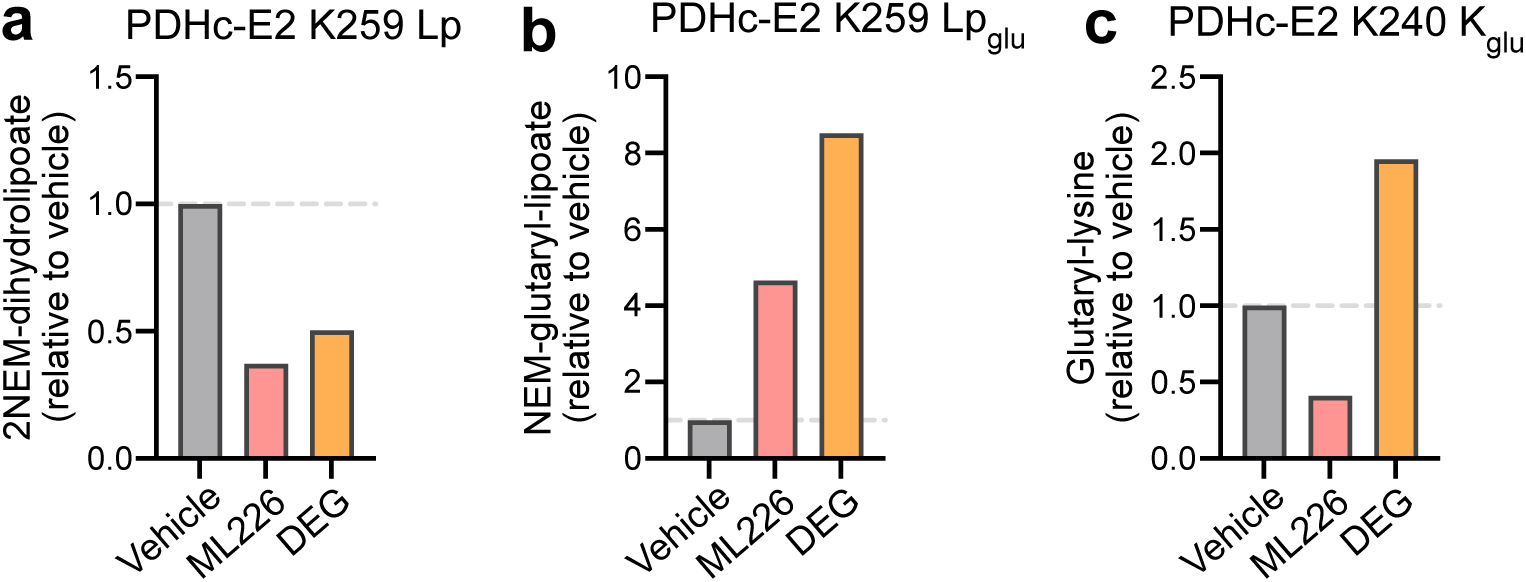
ABHD11 inhibition creates glutaryl-lipoate adducts on PDHc. **(a-c)** LC-MS/MS analysis of HeLa cells treated with 1 μM ML226 or 1 mM DEG for 24 hr, or 2.5 μM ML226 or 1 mM DEG for 6 hr, and PDHc-E2 was immunoprecipitated. Vehicle control and DEG treatment are the same as detailed by Minogue et al^6^. Quantification of PDHc-E2 K259 Lp **(a)**, K259 Lp_glu_ **(b)**, and K240 K_glu_ **(c)** by LC-MS/MS. Peptide abundance is normalised to the total abundance of modified peptides and adjusted relative to the vehicle condition.

**Extended Data Fig. 3.**
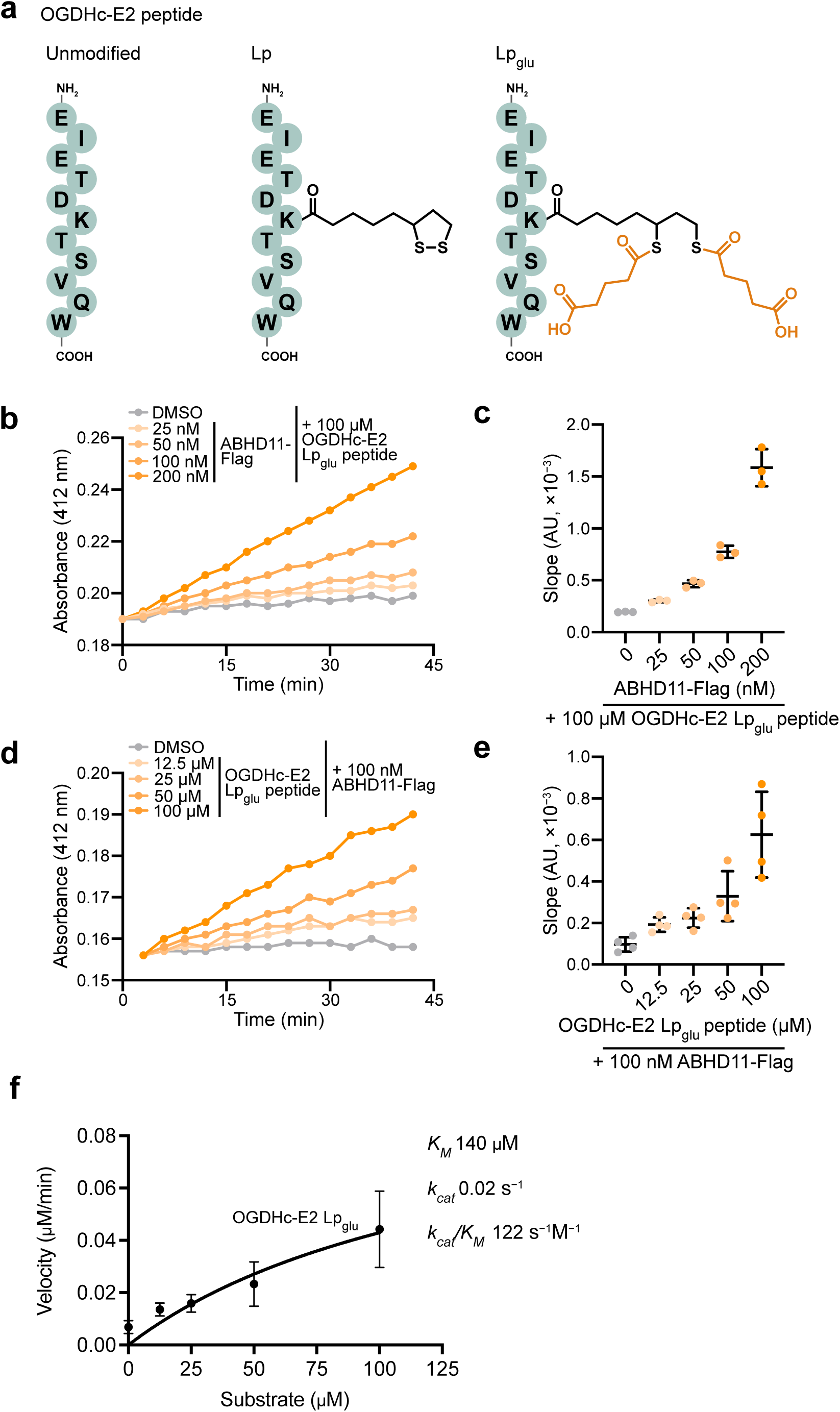
Glutaryl-lipoyl thioesterase activity of ABHD11. **(a)** Unmodified, Lp, and Lp_glu_ synthetic OGDHc-E2 peptides. **(b, c)** Thioesterase activity of ABHD11 against OGDHc-E2 Lp_glu_ (100 μM) at increasing enzyme concentrations. *n = 3; mean ± SD*. **(d, e)** ABHD11 activity (100 nM) against increasing concentrations of the OGDHc-E2 Lp_glu_ substrate. **(f)** Michaelis-Menten kinetics of ABHD11 thioesterase activity to OGDHc-E2 Lp_glu_.

**Extended Data Fig. 4.**
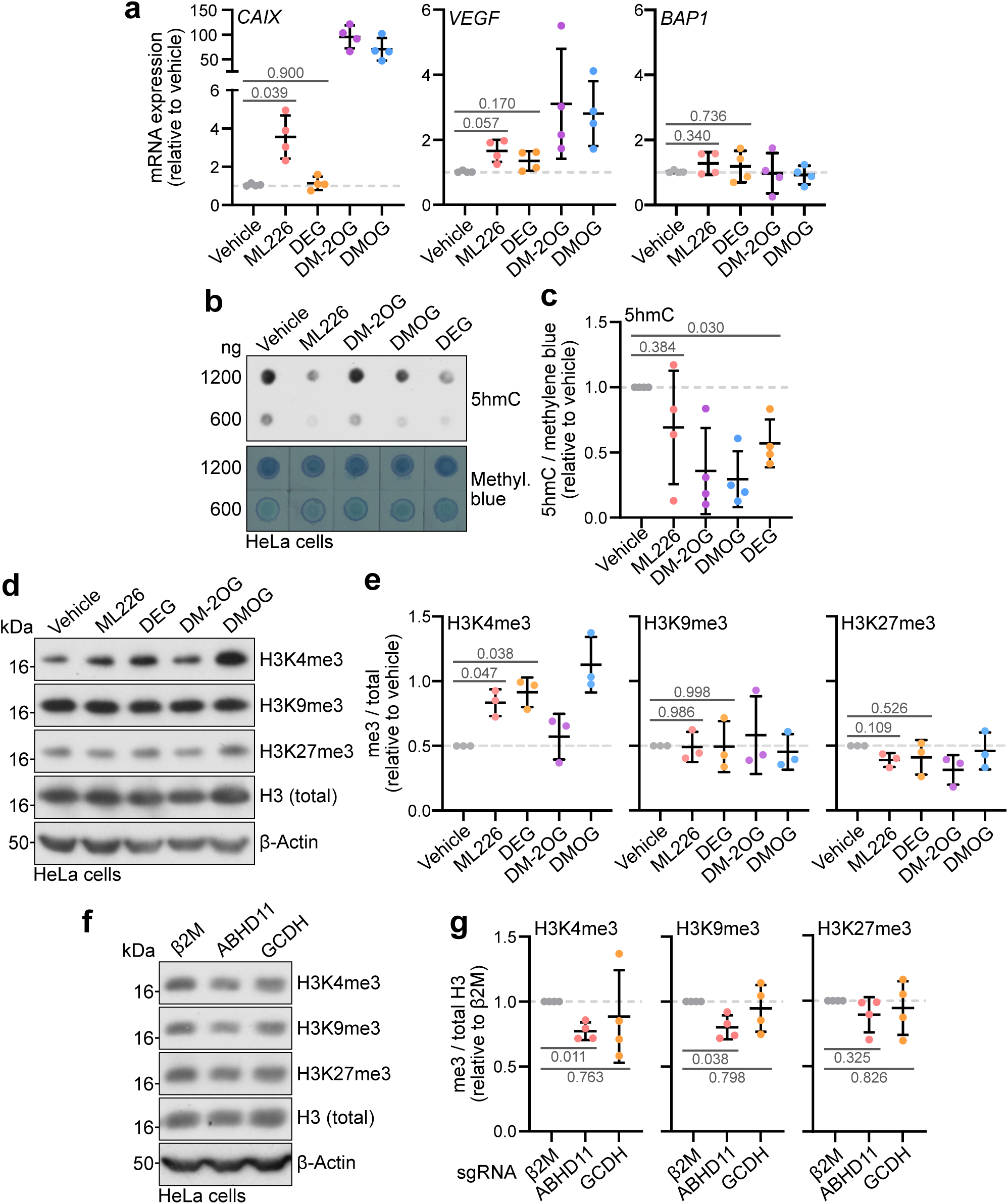
Glutaryl-lipoylation has distinct outcomes compared to protein glutarylation. **(a)** HeLa cells were treated with 1 μM ML226, 1 mM DEG, 1 mM DMOG or 6 mM DM-2OG for 24 h and *CAIX*, *VEGF*, and *BAP1* mRNA levels were determined using qRT-PCR. Transcript levels are normalised to *ACTB* and adjusted relative to the vehicle condition. *Mean ± SD; n = 4; one-way ANOVA and Dunnett’s post-hoc test.* **(b, c)** HeLa cells were treated with 2.5 μM ML226, 1 mM DEG, 1 mM DMOG or 6 mM DM-2OG for 24 h and 5hmC levels were determined using dot blotting **(b)**. Quantification of 5hmC levels **(c)** normalised to total DNA staining by methylene blue and adjusted relative to the vehicle condition. *Mean ± SD; n = 4; one-way ANOVA and Dunnett’s post-hoc test.* **(d, e)** HeLa cells were treated as in **(b)** and H3K4me3, H3K9me3, and H3K27me3 levels were determined using immunoblotting. Quantification of histone methylation levels **(e)** normalised to total histone H3 levels and adjusted relative to the vehicle condition. *Mean ± SD; n = 3; one-way ANOVA and Dunnett’s post-hoc test.* **(f, g)** Cas9-expressing HeLa cells were transduced with sgRNAs targeting ABHD11 or GCDH for 11 days and H3K4me3, H3K9me, and H3K27me3 levels were determined using immunoblotting. β2M sgRNA was used as a control. Quantification of histone methylation marks **(g)** normalised to total histone H3 levels and adjusted relative to the β2M condition. *Mean ± SD; n = 4; one-way ANOVA and Dunnett’s post-hoc test*.

**Extended Data Fig. 5.**
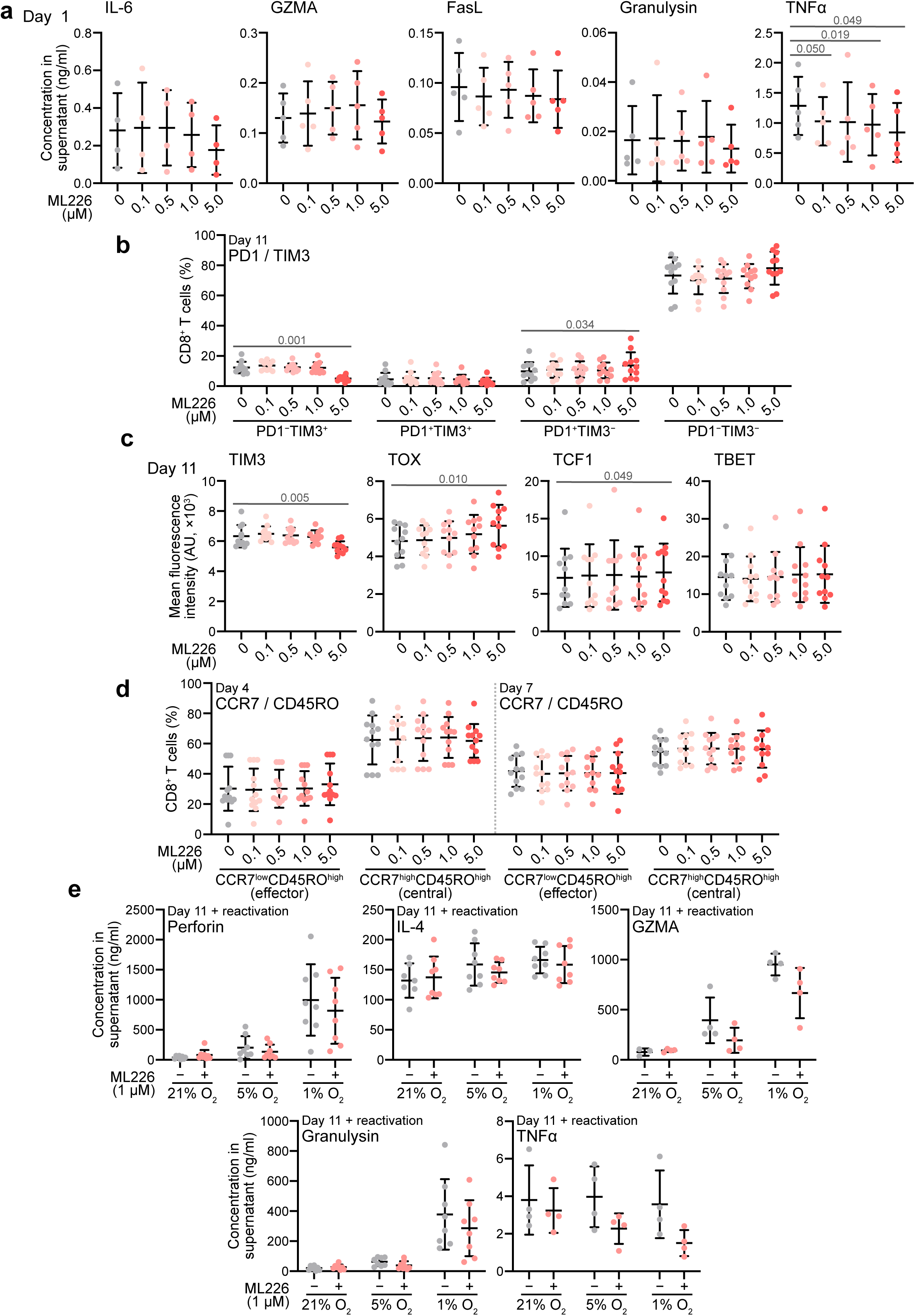
ABHD11 inhibition influences CD8+ T cell differentiation. **(a)** Cytokine levels in cell culture supernatants human CD8+ T cell activation assays (Fig. 5d). Activation markers 1 day post-activation were determined using flow cytometry. Each data point represents one human donor. *Mean ± SD; n = 5; one-way ANOVA and Dunnett’s post-hoc test.* **(b)** Percentage of PD1 and TIM3-expressing cells in **(a)** 11 days post-activation were determined using flow cytometry. *Mean ± SD; n = 11; one-way ANOVA and Dunnett’s post-hoc test for each subpopulation.* **(c)** Transcription factor (TIM3, TOX, TCF1, TBET) levels in **(a)** 11 days post-activation were determined using flow cytometry. *Mean ± SD; n = 11; one-way ANOVA and Dunnett’s post-hoc test.* **(d)** Percentage of CCR7 and CD45RO-expressing cells in **(a)** 4 or 7 days post-activation were determined using flow cytometry. *Mean ± SD; n = 12; one-way ANOVA and Dunnett’s post-hoc test for each timepoint.* **(e)** Cytokine levels in cell culture supernatants from hypoxia experiments (Fig. 5i) were determined using flow cytometry. Mean ± SD; n = 4–8; *Two-way ANOVA and Dunnett’s post-hoc test*.

**Extended Data Fig. 6.**
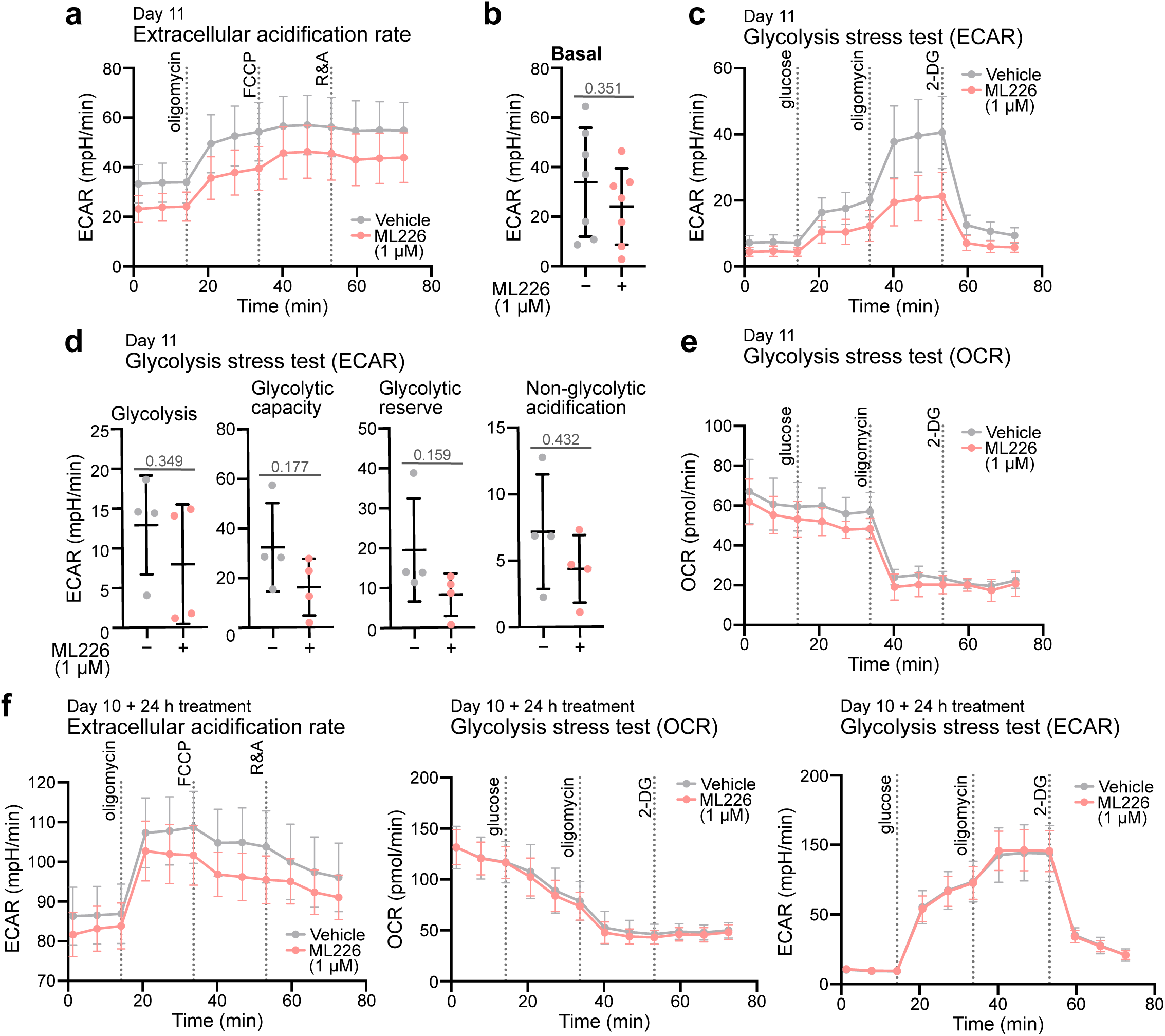
Effect of ABHD11 inhibition on CD8+ T cell metabolism. **(a, b)** CD8+ T cells were isolated from healthy donor peripheral blood mononuclear cells, activated with CD3/CD28 beads, and continuously treated with 1 mM ML226 for 11 days. Extracellular acidification (ECAR) during a mitochondrial stress test was quantified using a Seahorse XFe96 Extracellular Flux Analyzer. Quantification of basal ECAR **(b)**. *Mean ± SEM **(a)** or SD **(b)**; n = 6; unpaired two-tailed t-test.* **(c, d)** Activated CD8^+^ T cells were treated as in **(a)** and ECAR during a glycolysis stress test (10 mM glucose, 1 μM oligomycin, and 50 mM 2-deoxy-D-glucose (2-DG) was quantified using a Seahorse XFe96 Extracellular Flux Analyzer (mean ± SEM; n = 4). Quantification of glycolysis, glycolytic capacity, glycolytic reserve, and non-glycolytic acidificiation **(d)**. *Mean ± SEM **(c)** or SD (**d)**; n = 4; unpaired two-tailed t-test.* **(e)** Activated CD8+ T cells were treated as in **(a)** and OCR during a glycolysis stress test was quantified using a Seahorse XFe96 Extracellular Flux Analyzer. *Mean ± SEM; n = 4.* **(f)** Activated CD8+ T cells were cultured for 10 days and treated with 1 mM ML226 for 24 hours. ECAR during a mitochondrial stress, OCR during a glycolysis stress, and ECAR during a glycolysis stress was quantified using a Seahorse XFe96 Extracellular Flux Analyzer. *Mean ± SEM; n = 4*.

