## Supplementary Table 1 for "ABHD11 mediated deglutarylation regulates the TCA cycle and T cell metabolism"

**Supplementary Data 1**

**Reagents**

| **Reagent** | **Catalogue #** | **Source** |
| --- | --- | --- |
| ML226 | Cat#25681-1mg-CAY | Cayman Chemicals |
| Diethylglutarate (DEG) | QB-1473 | Combi-block |
| 5,5′-dithiobis-(2-nitrobenzoic acid) (DTNB) | D8130-1G | Sigma |
| Glutaryl CoA (Glutaryl-CoA Salt) | G9510-5MG | Sigma |
| Dimethyl 2-oxoglutarate  (DM-2OG) | 349631-5G | Sigma |
| Dimethyloxalylglycine (DMOG) | 71210-100 mg-CAY | Cambridge Bioscience |
| N-ethylmaleimide (NEM) | E3876-5G | Sigma |
| Nicotinamide | 72340-100G | Sigma |
| Puromycin | 100552-25MG | MP Biomedicals |

**SgRNA sequences**

| **Target** | **sgRNA sequence** |
| --- | --- |
| β2M | GGCCGAGATGTCTCGCTCCG |
| ABHD11 | AAGATCTTGGCCCAGCAGAC |
| GCDH (1)  GCDH (2)  GCDH (3) | GAGGCCGTGAACACCTACGA  CAGCAGAGAAACCATCTCGG  GATAGGGTGCATGACGAGGG |

**Antibodies**

| **Antibodies for immunoprecipitation and immunoblotting** | | |
| --- | --- | --- |
| **Antibody** | **Catalogue #, Source** | **Working dilution** |
| 5-hydroxymethylcytosine (rabbit serum) | Active Motif 39769 | 1:10,000 in 2% BSA in PBST |
| ABHD11  (rabbit polyclonal) | Enogene E11-14208C | 1:2,000 in 2% BSA in PBST |
| DLAT (4A4-B6-C10)  (mouse monoclonal) | Cell Signaling Technology 12362 | 1:2,000 in 2% BSA in PBST |
| DLST (9F4BD5) (mouse monoclonal) | Abcam ab110306 | 10μl for 5x10^7^ cells |
| DLST (D22B1) (rabbit monoclonal) | Cell Signaling Technology 11954 | 1:2,000 in 2% BSA in PBST |
| GCDH  (rabbit polyclonal) | Proteintech 14930-1-AP | 1:2,000 in 5% skim milk in PBST |
| Glutaryl-lysine  (rabbit polyclonal) | PTM Bio PTM-1151 | 1:2,000 in 5% skim milk in PBST |
| H3 (D1H2)  (rabbit monoclonal) | Cell Signaling Technology 4499 | 1:2,000 in 2% BSA in PBST |
| H3K27me3 (C36B11)  (rabbit monoclonal) | Cell Signaling Technology 9733 | 1:1,000 in 2% BSA in PBST |
| H3K4me3 (C42D8)  (rabbit monoclonal) | Cell Signaling Technology 9751 | 1:1,000 in 2% BSA in PBST |
| H3K9me3 (D4W1U)  (rabbit monoclonal) | Cell Signaling Technology 13969 | 1:1,000 in 2% BSA in PBST |
| HIF1α (D1S7W)  (rabbit monoclonal) | Cell Signaling Technology 36169 | 1:2,000 in 2% BSA in PBST |
| Lipoic acid  (rabbit polyclonal) | Sigma Aldrich 437695 | 1:2,000 in 2% fatty acid-free BSA in PBST |
| β-Actin  (mouse monoclonal) | Sigma-Aldrich A2228 | 1:20,000 in 2% BSA in PBST |
| **Flow cytometry antibodies** | | |
| **Antibody** | **Catalogue #** | **Source** |
| CD8 (RPA-T8) | BUV395 | BD Biosciences 563796 |
| CD45RO (UCHL1) | BUV495 | BD Biosciences 749888 |
| CD25 (BC96) | BV510 | Biolegend 302639 |
| CD45RA (HI101) | BV650 | Biolegend 304135 |
| CD62L (DREG-56) | AF488 | Biolegend 304816 |
| CD62L (DREG-56) | PerCP/Cy5.5 | Biolegend 304824 |
| CCR7 (3D12) | PE/Cy7 | BD Biosciences 560922 |
| CD27 (LG.3A10) | AF700 | Biolegend 124239 |
| TIM3 (F38-2E2) | BV605 | Biolegend 345017 |
| PD1 (EH12.2H7) | AF488 | Biolegend 329936 |
| TCF1 (S33-966) | PE | BD Biosciences 564217 |
| LAG3 (11C3C65) | AF647 | Biolegend 369304 |
| Perforin (dG9) | Pacific Blue | Biolegend 308117 |
| Granzyme B (AD2) | PerCP/Cy5.5 | Biolegend 344013 |
| TBET (4B10) | PE/Dazzle594 | Biolegend 644828 |
| TIGIT (A15153G) | PE/Cy7 | Biolegend 372713 |
| TOX (TXRX10) | eflour660 | eBiosciences 50-6502-82 |

**Quantitative PCR primers**

| **Target gene** | **Forward primer** | **Reverse primer** |
| --- | --- | --- |
| ACTB | CTGGGAGTGGGTGGAGGC | TCAACTGGTCTCAAGTCAGTG |
| CAIX | GCCGCCTTTCTGGAGGA | TCTTCCAAGCGAGACAGCAA |
| VEGF | TACCTCCACCATGCCAAGTG | ATGATTCTGCCCTCCTCCTTC |
| BAP1 | GGAGGTAGAGAAGAGGAAGAA | GAGCCAGCATGGAGATAAAG |

**Software**

| **Software** | **Source** | **Identifier** |
| --- | --- | --- |
| FlowJo v10 | BD Biosciences | RRID:SCR_008520 |
| Peaks 11 | Bioinfor | https://www.bioinfor.com/peaks-11 |
| Prism v9.5.1 | GraphPad | RRID:SCR_002798 |
| ProMod3 3.3.0 | swissmodel.expasy.org |  |
| PyMOL 3.0.4 | Pymol | RRID:SCR_000305 |
| Coot | https://www2.mrc-lmb.cam.ac.uk/personal/pemsley/coot/ | RRID:SCR_014222 |
| SHELX | http://shelx.uni-ac.gwdg.de/SHELX/ | RRID:SCR_014220 |
| HADDOCK2.4 | https://rascar.science.uu.nl/haddock2.4/ | RRID:SCR_019091 |
